# The CD8^+^ T cell landscape of human brain metastases

**DOI:** 10.1101/2021.08.03.455000

**Authors:** Lisa J. Sudmeier, Kimberly B. Hoang, Edjah K. Nduom, Andreas Wieland, Stewart G. Neill, Matthew J. Schniederjan, Suresh S. Ramalingam, Jeffrey J. Olson, Rafi Ahmed, William H. Hudson

**Affiliations:** Department of Radiation Oncology, Emory University School of Medicine, Atlanta, GA USA; Winship Cancer Institute, Emory University School of Medicine, Atlanta, GA USA; Department of Neurological Surgery, Emory University School of Medicine, Atlanta, GA USA; Emory Vaccine Center, Emory University School of Medicine, Atlanta, GA USA; Department of Microbiology and Immunology, Emory University School of Medicine, Atlanta, GA USA; Department of Pathology and Laboratory Medicine, Emory University School of Medicine, Atlanta, GA USA; Department of Hematology and Medical Oncology, Emory University School of Medicine, Atlanta, GA USA

## Abstract

Despite improved outcomes with checkpoint blockade immunotherapy, patients with brain metastases have the worst prognosis among patients with metastatic cancer. Immune checkpoint blockade agents target inhibitory receptors, such as PD-1, on exhausted CD8^+^ T cells to restore their anti-cancer function. Many patients, however, either do not respond or progress after an initial response to immune checkpoint blockade, and distant intracranial failure is common despite excellent options for local treatment of brain metastasis. To develop more effective therapeutic strategies for the treatment of brain metastases, an understanding of the phenotype of brain metastasis-infiltrating CD8^+^ T cells is essential. Here we performed a detailed characterization of the CD8^+^ T cells contained in brain metastases. Brain metastases were densely infiltrated by CD8^+^ T cells; blood contamination of tumor samples was rare. Compared to patient-matched circulating cells, brain metastasis-infiltrating CD8^+^ T cells had a distinct phenotype characterized by more frequent expression of PD-1, with subpopulations defined by expression of additional co-inhibitory molecules and the residence marker CD69. Single cell RNA-sequencing identified four phenotypic subpopulations within brain metastasis-infiltrating PD-1^+^ CD8^+^ T cells. Two of these populations — a terminally-differentiated and a dividing population — were characterized by high expression of co-inhibitory molecules and lacked expression of progenitor markers such as TCF-1. There was significant T cell receptor (TCR) overlap between the terminally-differentiated and dividing populations, suggesting that the dividing cells give rise to the terminally-differentiated cells. There was minimal TCR overlap between these two populations and other brain metastasis-infiltrating PD-1^+^ CD8^+^ T cells. T cell clones from brain metastasis-infiltrating CD8^+^ T cells were rare in circulation, particularly clones from the terminally-differentiated and dividing populations. We systematically identified bystander CD8^+^ T cells specific for microbial antigens; these cells infiltrated brain metastases and expressed genes shared with exhausted progenitor CD8^+^ T cells, such as *TCF7* and *IL7R*. We performed spatial transcriptomics on brain metastases and used a novel method to obtain TCR sequences from spatial transcriptomics data. These data revealed distinct niches within the TME defined by their gene expression patterns and cytokine profiles. Terminally-differentiated CD8^+^ T cells preferentially occupied niches within the tumor parenchyma. Together, our results show that antigen-specificity restricts the spatial localization, phenotypic states, and differentiation pathways available to CD8^+^ T cells within the brain metastasis TME.

## Introduction

Brain metastases affect approximately 20% of all cancer patients^1^ and are associated with significantly worse overall survival compared to extracranial metastases.^2^ Although they respond to immune checkpoint blockade (ICB),^3^ brain metastases also receive local therapy - radiation therapy with or without surgical resection^4^ - so the isolated effect of CD8^+^ T cell reinvigoration after ICB on these tumors has been difficult to discern. The brain comprises a unique immune environment, classically thought to be immunosuppressive, which protects the central nervous system (CNS) from excessive inflammation.^5^ Although there is loss of blood-brain-barrier (BBB) integrity within brain tumors,^6,7^ the consequent less-selective blood-tumor-barrier (BTB) and surrounding stroma of brain tissue may together affect inflammatory signaling and cell recruitment to the tumor microenvironment (TME) of brain metastases. Brain metastases are infiltrated by CD8^+^ T cells^8–12^ which may not be clonally related to CD8^+^ T cells infiltrating patient-matched primary tumors^13^, suggesting that brain metastases may have different determinants governing response to ICB compared to the primary tumor.

CD8^+^ T cells are cytotoxic lymphocytes that recognize infected or neoplastic cells via antigen-specific binding of their T cell receptor (TCR) to MHC class I-peptide complexes. Upon antigen clearance, effector CD8^+^ T cells differentiate into memory cells which protect against antigen re-exposure.^14^ In cases where antigen is not cleared, as in cancer or chronic infections, effector CD8^+^ T cells become exhausted, a state characterized by impaired proliferative and cytotoxic capacity.^15^ Exhausted CD8^+^ T cells express inhibitory molecules such as PD-1, which further promote the exhausted phenotype^16^. PD-1 and other inhibitory pathways are the targets of ICB agents, which block suppressive signaling in exhausted CD8^+^ T cells to rescue their proliferative and cytotoxic function^17^. Clinical use of ICB agents in the treatment of cancer has resulted in dramatic improvements in disease control and patient survival, however a significant proportion of cancer patients have refractory disease that either does not respond to ICB or progresses after an initial response^18,19^. One strategy to improve ICB efficacy is to simultaneously block multiple inhibitory molecules expressed on exhausted CD8^+^ T cells^20^. In order to identify potential therapeutic targets for this combination approach, a detailed phenotypic characterization of the target exhausted CD8^+^ T cells is required.

Exhausted CD8^+^ T cells are comprised of diverse phenotypic subpopulations with distinct functions, inhibitory molecule expression, and tissue homing patterns. Exhausted progenitor PD-1^+^ CD8^+^ T cells, which are maintained by expression of the transcription factor TCF-1, have the capacity to self-renew and produce daughter cells that undergo further differentiation ^21–23^. Transitory PD-1^+^ CD8^+^ T cells are the immediate progeny of these progenitor cells and are characterized by expression of effector molecules and loss of TCF-1. These transitory cells are migratory; circulating antigen-specific CD8^+^ T cells in cancer and chronic infection are found in this state, which is marked by CX3CR1 expression.^24–29^ Upon migration to non-lymphoid tissues, these transitory cells further differentiate into a terminally-differentiated population with increased expression of inhibitory molecules.^24–26^ These most-exhausted CD8^+^ T cells are resident in non-lymphoid tissues, have poor effector function, and lack proliferative capacity.^24,26^

These CD8^+^ T cell populations each respond differently to PD-1 pathway blockade. The exhausted progenitor population is required for the proliferative burst observed after ICB, which produces significant expansion in the number of transitory effector cells^21–23^. ICB also acts on effector CD8^+^ T cells at the site of antigen to improve their effector function by increasing their expression of molecules such as granzymes and perforins.^23,24,30,31^ Given its requirement for the proliferative burst in response to PD-1 pathway blockade, the progenitor population of exhausted CD8^+^ T cells has received significant attention in tumor immunology studies, many of which have quantified tumor-infiltrating TCF-1^+^ cells^26,32–34^. However, the antigen specificity of tumor-infiltrating TCF-1^+^ CD8^+^ T cells is rarely determined.

Tumor-specific exhausted progenitor TCF-1^+^CD8^+^ T cells have been found in human papillomavirus positive (HPV+) head and neck squamous cell carcinoma^35^, which grows in oropharyngeal lymphoid tissues.^36^ Similar progenitor cells were also recently identified in melanoma and non-small cell lung cancers, where a majority of tumor-specific CD8^+^ T were in a terminally differentiated state^37,38^. In murine models, antigen-specific exhausted progenitor CD8^+^ T cells are enriched in tumor-draining lymph nodes and are found exclusively in secondary lymphoid organs during chronic infection^39–41^. However, it is unclear whether tumor-specific exhausted progenitor CD8^+^ T cells reside in tumors growing in the brain. In the absence of antigen-specificity information, CD8^+^ T cell function is often inferred from phenotype. However, this approach is confounded by the expression of some molecular markers at multiple stages of CD8^+^ T cell differentiation. TCF-1, for example, is expressed by naïve, memory, and exhausted progenitor CD8^+^ T cells. CMV- and EBV-specific effector memory CD8^+^ T cells co-express TCF-1 and TOX,^42^ a transcription factor associated with T cell exhaustion,^43–45^ and EBV- and influenza-specific CD8^+^ T cells have been found in primary and metastatic brain tumors.^46^ Moreover, brain-resident memory cells have been reported to maintain antigen-independent PD-1 expression.^47^

Here we performed a detailed characterization of brain metastasis-infiltrating CD8^+^ T cells and their surrounding TME from a cohort of 31 patients who underwent surgical resection of a large and/or symptomatic brain metastasis. Brain metastases were well-infiltrated by CD8^+^ T cells, a majority of which were PD-1^+^. Using single cell RNA sequencing (scRNA-seq), we identified four transcriptional populations among PD-1^+^ CD8^+^ T cells infiltrating brain metastases: dividing cells, terminally-differentiated cells, and two clusters that shared some phenotypic features with exhausted progenitor cells. These first two subsets shared significant clonal overlap with each other, but had minimal T cell receptor (TCR) overlap with the progenitor-like populations. We systematically identified bystander cells specific for microbial antigens among brain metastasis-infiltrating CD8^+^ T cells; these were rare in the terminally-differentiated and dividing populations and had a phenotype similar to that of exhausted progenitor cells. Bystanders were present among brain metastasis-infiltrating PD-1^+^ CD8^+^ T cells and circulating PD-1^+^ CD8^+^ T cells at a similar frequency. To determine the location of specific CD8^+^ T cell clones within the tumor, we used a novel method to obtain TCR sequences from spatial transcriptomics data and showed that CD8^+^ T cells from each phenotypic population were spatially restricted to specific regions of the brain metastasis TME with distinct gene expression patterns and cytokine profiles. Together, our results show that brain metastases are infiltrated by diverse populations of CD8^+^ T cells that adopt specific phenotypes and segregate to distinct niches within the TME based on their antigen specificity. These data may guide novel immunotherapeutic strategies for the treatment of brain metastases.

## Results

### Human brain metastases are well-infiltrated by CD8^+^ T cells

Over 18 months we collected fresh brain metastasis tumor specimens and matched blood samples from 31 patients at Emory University Hospital who underwent surgical resection of at least one brain metastasis (Supplementary Table 1). Samples were obtained fresh at time of surgical resection and included a mixture of primary tumor types, the most abundant of which was lung carcinoma (Fig. 1a), consistent with it being the primary cancer most likely to metastasize to the brain.^2,48^ All patients were naïve to immunotherapy. The immune infiltrate of all samples was quantified by flow cytometry. A subset of samples was used for high-parameter flow cytometry, single-cell RNA sequencing (scRNA-seq), T cell receptor (TCR) sequencing, and immunohistochemistry with spatially-resolved transcriptomics (Fig. 1a).

**Figure 1:**
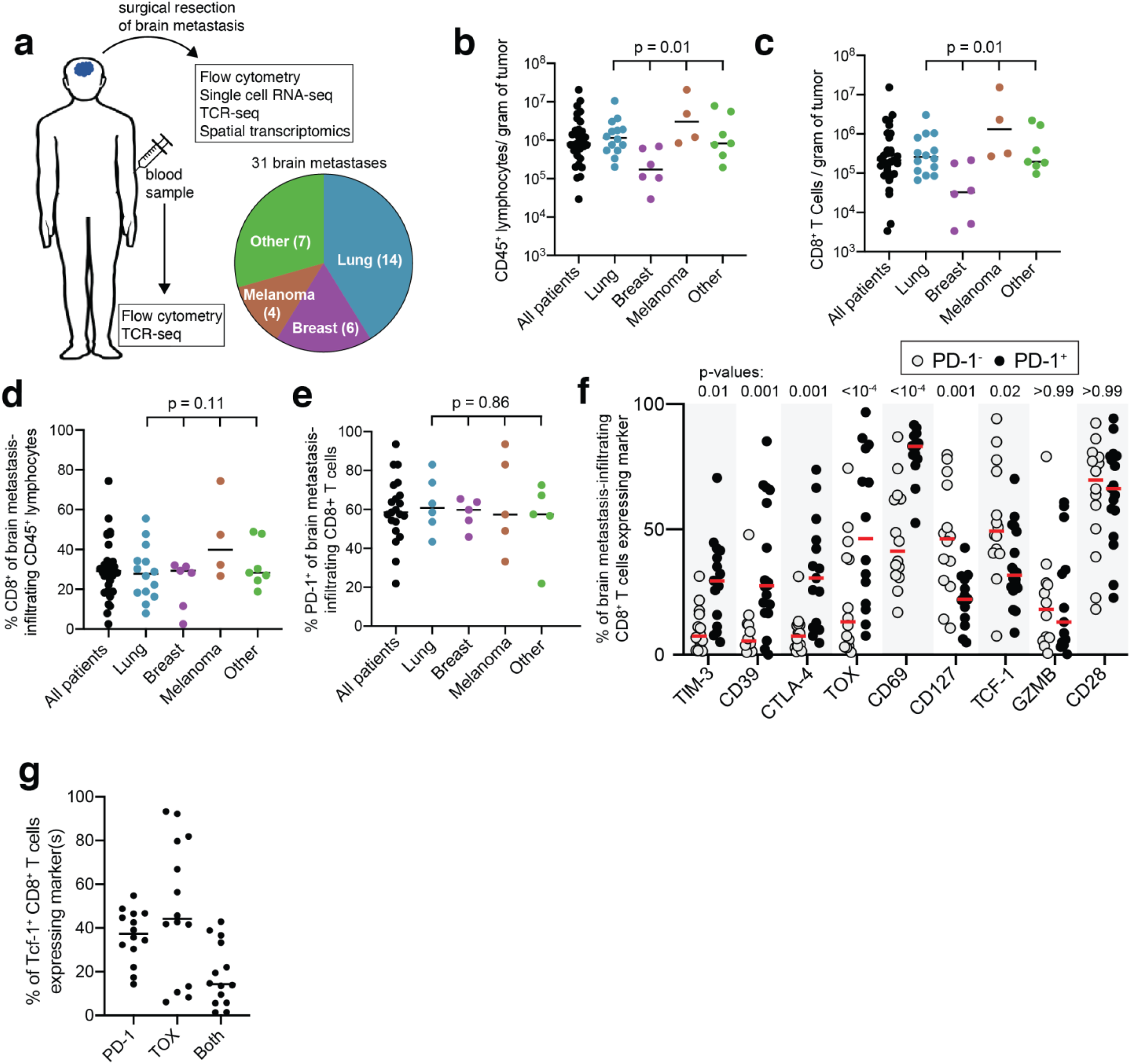
Human brain metastases are well-infiltrated by PD-1^+^ CD8^+^ T cells. (a) Experimental schema and distribution of samples among primary tumor histologies. (b) Density of CD45^+^ lymphocytes and (c) CD8^+^ T cells for all 31 tumors, grouped by primary tumor type. (d) Frequency of CD8^+^ lymphocytes grouped by tumor type. (e) Percentage of CD8^+^ T cells expressing PD-1. (f) Phenotype of PD-1^−^ vs. PD-1^+^ CD8^+^ T cells. (g) Percent of TCF-1^+^ CD8^+^ T cells expressing PD-1, TOX, or both. Bars on graphs indicate median. In panels (b) and (c) statistics show statistically significant variance among primary tumor types with the Kruskal-Wallis test. In panels (d) and (e) statistics show one-way ANOVA. In (f) statistics show a mixed-effects model analysis with Sidak’s multiple comparisons test.

Brain metastases were variably infiltrated by immune cells, ranging from 2.9 × 10^4^ to 2 × 10^7^ CD45^+^ lymphocytes per gram of tumor (median 7.6 × 10^5^), with brain metastases from breast cancer having the lowest infiltration density (Fig. 1b). CD8^+^ T cell infiltration was also variable (range 3.4 × 10^3^ to 1.5 × 10^7^, median 2.0 × 10^5^ CD8^+^ T cells/gram of tumor), with breast carcinoma histology again showing the lowest density of infiltration (Fig. 1c). Despite significant variability among samples, CD8^+^ T cells comprised a similar proportion of lymphocytes within brain metastases, regardless of primary tumor type (Fig. 1d).

### A subset of brain metastasis-infiltrating CD8^+^ T cells have a progenitor phenotype

To interrogate the phenotype of brain metastasis-infiltrating CD8^+^ T cells, we performed high-parameter flow cytometry on patient-matched tumor-infiltrating and circulating CD8^+^ T cells. The frequency of brain metastasis-infiltrating CD8^+^ T cells expressing PD-1 was consistent across different tumor histologies despite some variation (Fig. 1e). We compared the expression of other co-inhibitory molecules, transcription factors, and the effector molecule granzyme B (GZMB) on PD-1^−^ and PD-1^+^ CD8^+^ T cells (Fig. 1f, Supplementary Fig. 1). TOX and the co-inhibitory molecules TIM3, CD39 and CTLA-4 were significantly higher on PD-1^+^ cells compared to PD-1^−^ cells. Although CD69 was more highly expressed on PD-1^+^ cells, nearly half of PD-1^−^ cells also expressed CD69, indicating that a portion of these cells are also resident in the tumor (Fig. 1f). While the effector molecule GZMB and the co-stimulatory molecule CD28 are expressed similarly between PD-1^+^ and PD-1-CD8^+^ T cells, markers of progenitor function of CD8^+^ T cells such as CD127 and TCF-1 are higher in PD-1^−^ cells (Fig. 1f).

Little is known about the abundance and phenotype of TCF-1^+^ CD8^+^ T cells in brain metastases, and it is unclear whether tumor-specific exhausted progenitor cells reside in these tumors. CD127 and CD28 have been used as extracellular markers of TCF-1^+^ progenitor CD8^+^ T cells in chronic infections and cancer^23,32,41^. In brain metastasis-infiltrating CD8^+^ T cells, CD28 and CD127 were both more frequent on PD-1^+^ TCF-1^+^ compared to PD-1^+^ TCF-1^−^ cells (Supplementary Fig. 1b-c), but their expression did not completely recapitulate that of TCF-1. CD127 was mostly absent from PD-1^+^ TCF-1^−^ cells, but only expressed on half of PD-1^+^ TCF-1^+^ cells (Supplementary Fig. 1b-c). CD28 was a more sensitive marker of TCF-1 expression, and was found on over 75% of TCF-1^+^ cells, but lacked specificity, with expression on over 50% of TCF-1^−^ cells (Supplementary Fig. 1b-c). The transcription factor TOX is a marker and regulator of CD8^+^ T cell exhaustion and is expressed on antigen-specific CD8^+^ T cells in both cancer and chronic infection^35,43–45^. In our cohort, 44% of brain metastasis-infiltrating TCF-1^+^ CD8^+^ T cells expressed TOX, and 37% were PD-1^+^; 14% of TCF-1^+^ CD8^+^ T cells co-expressed PD-1 and TOX (Fig. 1g). Thus, despite high frequency of TCF-1^+^ CD8^+^ T cells within brain metastases, these cells are phenotypically diverse and TCF-1 expression alone may not be an adequate marker of tumor-specific CD8^+^ T cell progenitor function.

### Phenotypically distinct populations of CD8^+^ T cells infiltrate brain metastases

FlowSOM clustering of all tumor-infiltrating and matched circulating CD8^+^ T cells identified six populations (Fig. 2a, Supplementary Fig. 2). Clusters 1 and 2 were preferentially found in blood, and clusters 4 and 5 were exclusively tumor-infiltrating (Fig. 2b-c). Cluster 3 was present in high frequencies in blood and tumor, and cluster 6 was a rare population in both blood and tumor (Fig. 2b-c). Cluster 1 is comprised of naïve CD8^+^ T cells, characterized by expression of CCR7 and CD45RA (Fig. 2d-e, Supplementary Fig. 2b); its low frequency in the tumor is indicative of minimal blood contamination of brain metastasis specimens (Fig. 2c). Cells in cluster 2 were predominately CD45RA^+^ and expressed high levels of granzyme B (Fig. 2e, Supplementary Fig. 2b). Cluster 3 was present in the circulating and tumor compartments and was comprised of heterogenous PD-1^−^ and PD-1^dim^ cells with high levels of CD28, CD127, and TCF-1 (Fig. 2c-e). Cells in clusters 4 and 5 both expressed CD69, consistent with tissue residence and their predominance in the tumor (Fig. 2e, Supplementary Fig. 2b).

**Figure 2:**
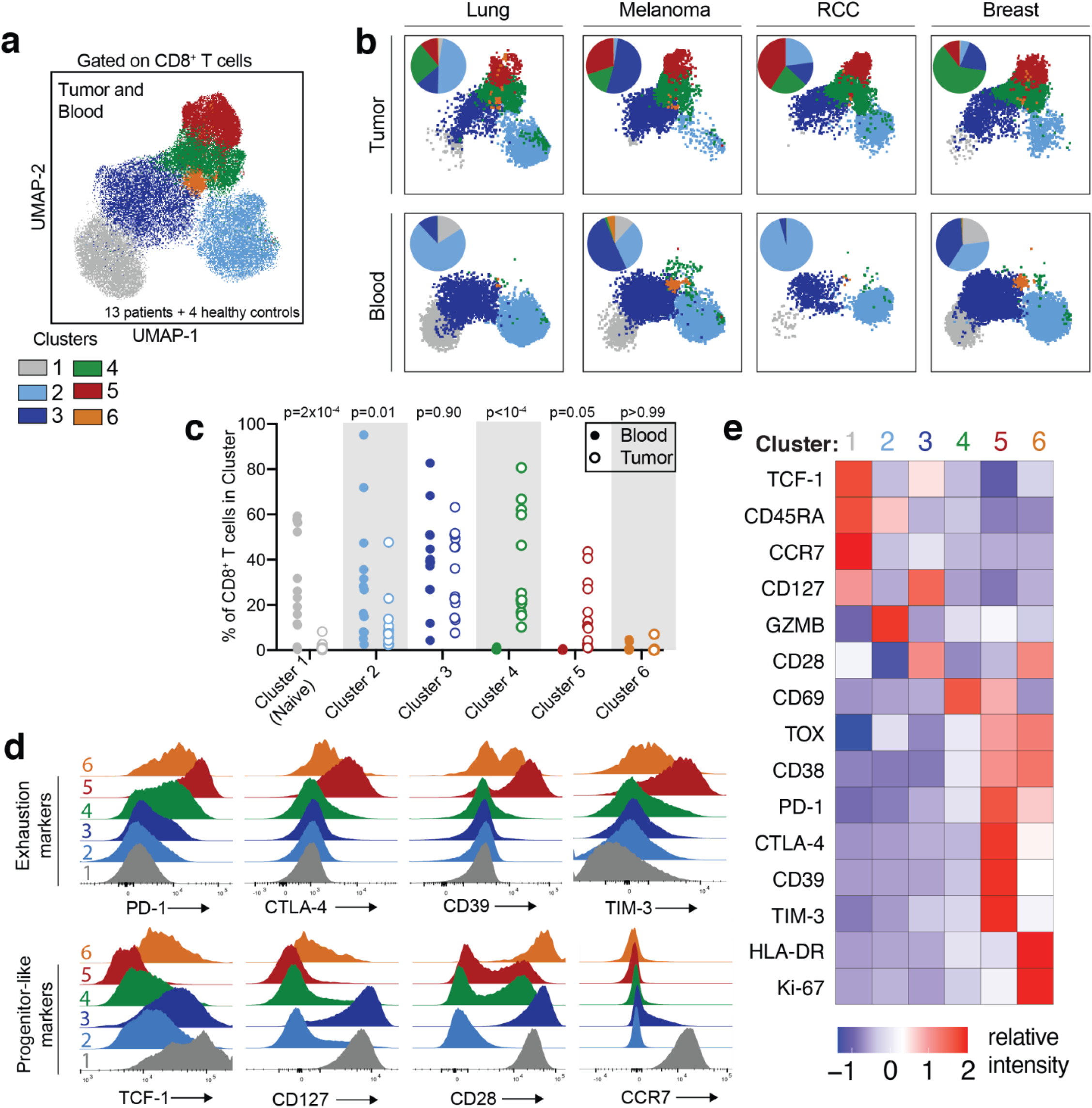
Spectral flow cytometry reveals that brain metastases are infiltrated by CD8^+^ T cells that are phenotypically distinct from circulating CD8^+^ T cells. (a) UMAP projection of high parameter flow cytometry data gated on CD8^+^ T cells from 13 brain metastases, patient-matched blood, and blood from four healthy controls. Cells are colored by FlowSOM cluster. (b) UMAPs of CD8^+^ T cells from four individual patients (RCC, renal cell carcinoma). Tumor-infiltrating and circulating CD8^+^ T cells are shown in the top and bottom row, respectively. Inlaid pie charts show percentage of cells in each FlowSOM cluster. (c) Percent of CD8^+^ T cells in each cluster is shown for each patient’s blood and tumor sample. (d) Expression of selected markers in each CD8^+^ T cell cluster. (e) Relative intensity of each indicated marker within each flow cluster. Statistics in panel (c) are two-way ANOVA with Sidak’s multiple comparisons test.

Cells in both clusters 4 and 5 also expressed CD38 and were low for TCF-1 and CD127; expression of these latter two markers on CD8^+^ T cells within the tumor was primarily restricted to cluster 3, a cluster shared between the tumor and circulating compartments (Fig. 2c-e, Supplementary Fig. 2b). Cluster 5 appears to be an exhausted, terminally-differentiated population of CD8^+^ T cells, expressing PD-1 as well as other co-inhibitory molecules including CTLA-4, CD39 and TIM-3 (Fig. 2d-e). Most cells in cluster 5 also expressed TOX and were TCF-1^−^ (Fig. 2e, Supplementary Fig. 2b). Cells in cluster 6 are dividing (KI-67^+^) and express activation markers such as HLA-DR and CD38, as well as exhaustion markers such as PD-1 (Fig. 2e, Supplementary Fig. 2b). Cluster 6 cells express CD28 but low levels of TCF-1 and CD127 (Fig. 2d-e).

### Brain metastasis-infiltrating CD8^+^ T cells comprise four metaclusters with distinct transcriptional phenotypes

CD8^+^ T cells specific for tumor-associated viral and neoantigen epitopes express PD-1^35,49–51^. In our cohort, PD-1 expression within the tumor was highest on the terminally-differentiated cluster 5 and dividing cells (FlowSOM cluster 6; Fig. 2d-e). PD-1 was also expressed at lower levels in tumor-enriched cluster 4 and on some cells in cluster 3, which was shared between blood and tumor (Fig. 2c-e). To determine transcriptional profiles, interrogate differentiation pathways, and examine antigen-specificity of these PD-1-expressing cells, we performed scRNA-seq with TCR sequencing on sorted PD-1^+^ CD8^+^ T cells isolated from three non-small cell lung carcinoma (NSCLC) brain metastases and two melanoma brain metastases immediately after surgical resection (Supplementary Fig. 3). These two histologies were chosen because they commonly metastasize to the brain. Naïve CD8^+^ T cells were also sorted from patient-matched blood as a control. From 22,828 sequenced cells, we identified fourteen populations of PD-1^+^ CD8^+^ T cells, which were hierarchically clustered into 5 metaclusters with similar gene expression patterns: A, B, C, Dividing (D), and Naïve (Fig. 3a, Supplementary Table 2). Consistent with our flow cytometric analyses, naïve cells were found predominantly in blood. Four of the five patients had cells in each of clusters A-D, the exception being patient 17, who did not have cells in metacluster C (Fig. 3b). There was no difference in the percentage of cells in each metacluster between brain metastases from lung cancer vs melanoma, suggesting that tissue of origin did not strongly influence the phenotype of brain metastasis-infiltrating PD-1^+^ CD8^+^ T cells in our cohort (Fig. 3c-d).

**Figure 3:**
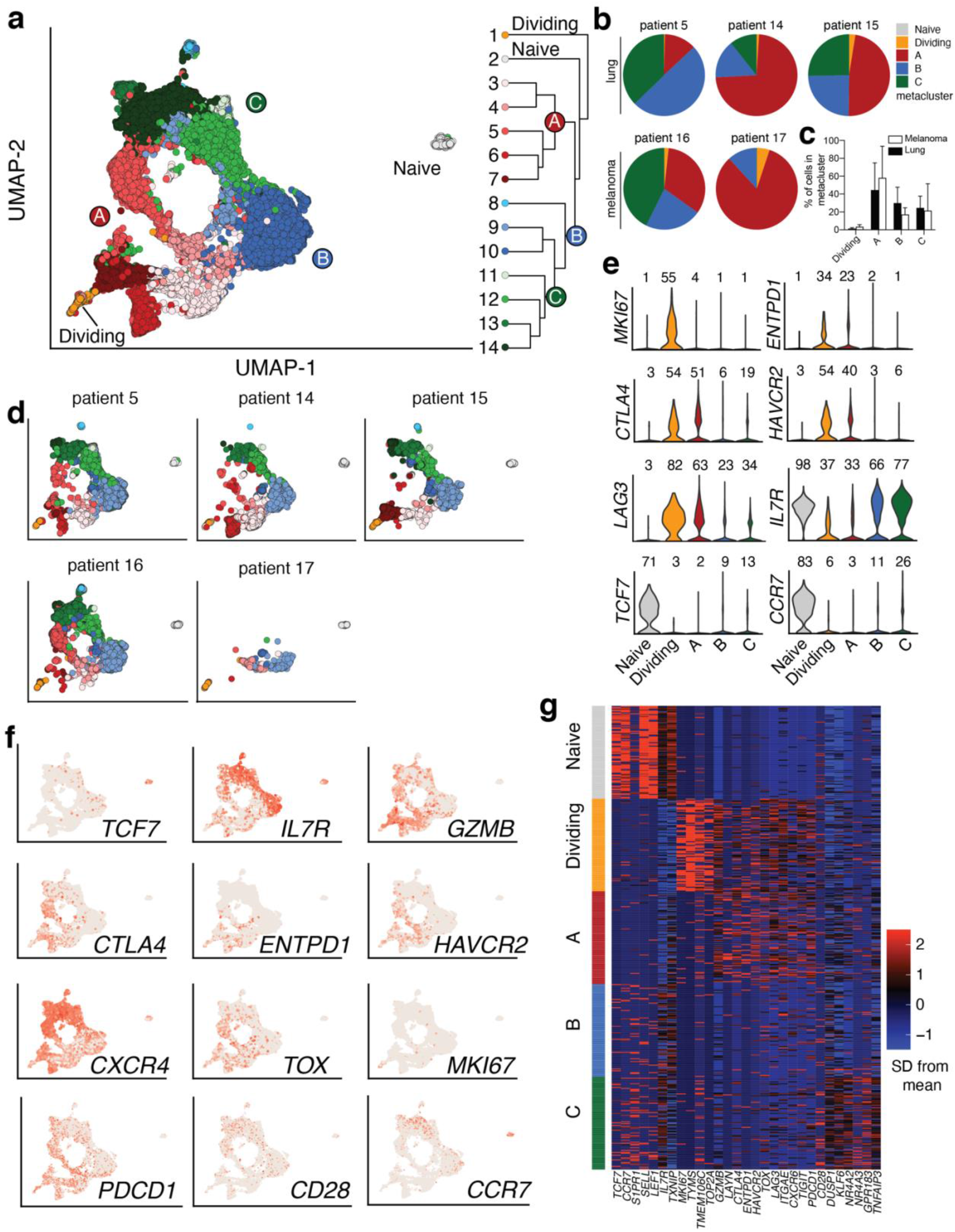
Transitory and terminally differentiated PD-1^+^ CD8^+^ T cells infiltrate human brain metastases. (a) UMAP of all 22,828 PD-1^+^ CD8^+^ T cells sorted from 5 brain metastases and naïve cells sorted from patient-matched blood samples. Hierarchical clustering is shown at right. (b) Distribution of each patient’s PD-1^+^ CD8^+^ T cells among metaclusters. (c) Distribution of PD-1^+^ CD8^+^ T cells among metaclusters for each primary tumor type. (d) Phenotype of PD-1^+^ CD8^+^ T cells from brain metastasis and matched circulating naïve cells for each patient. (e) Expression of selected genes in each metacluster. Numbers above each violin indicate percent of cells in each metacluster expressing the gene. (f) Expression of selected genes projected on UMAP. (g) Relative expression of selected genes in each metacluster.

Tumor-infiltrating PD-1^+^ CD8^+^ T cells in metaclusters A and D had a terminally-differentiated phenotype similar to that of FlowSOM cluster 5, characterized by high expression of genes encoding co-inhibitory molecules such as *CTLA4, ENTPD1* (CD39), *HAVCR2* (TIM-3), and *LAG3*, consistent with this population containing tumor-reactive cells (Fig. 3e-f, Supplementary Fig. 4). Cells in the Dividing metacluster were additionally defined by high expression of cell cycle genes, such as *MKI67* (KI-67) and *TOP2A* (Fig. 3g). Gene set enrichment analysis (GSEA) revealed that the transitory transcriptional signature characterized in the murine LCMV (lymphocytic choriomeningitis virus) model of CD8^+^ T cell exhaustion^24^ was enriched in the Dividing metacluster, suggesting that these cells may be undergoing differentiation from stem-like to terminally-differentiated PD-1^+^ CD8^+^ T cells (Supplementary Fig. 4). Metaclusters B and C expressed the lowest levels of co-inhibitory markers (Fig. 3e-f, Supplementary Fig. 4b). They were distinguished from each other by higher expression of tissue residence genes, such as CD69, in metacluster C (Supplementary Fig. 4a). Together, metaclusters B and C contained PD-1^+^ CD8^+^ T cells with higher expression of the progenitor markers *TCF7* (TCF-1) and *IL7R* (CD127) compared to terminally-differentiated and dividing cells. (Fig. 3e-g).

### Terminally-differentiated CD8^+^ T cells have minimal clonal overlap with circulating or progenitor-like tumor-infiltrating CD8^+^ T cells

To determine the clonal relationship between circulating and brain metastasis-infiltrating CD8^+^ T cells, we performed TCR sequencing on non-naïve PD-1^+^ and PD-1^−^ CD8^+^ T cells sorted from patient-matched peripheral blood (Supplementary Fig. 3). Compared to circulating PD-1^−^ CD8^+^ T cells, circulating PD-1^+^ CD8^+^ T cells had lower TCR diversity and more overlap with tumor-infiltrating cells (Fig. 4a). However, the overall overlap between circulating and tumor-infiltrating CD8^+^ T cells was minimal, suggesting that circulating CD8^+^ T cells expressing tumor-enriched TCRs are rare (Fig. 4b)^29,52^. Circulating tumor-specific CD8^+^ T cells appear to be even more infrequent, as TCRs from terminally-differentiated cells were rarely found in blood (Fig. 4c-d). Tumor-infiltrating cells that did express blood-enriched TCRs were predominantly located in metaclusters B and C (Fig. 4c-d). This is consistent with our flow cytometry data, where TCF-1 and CD127 expression in the tumor was restricted to FlowSOM cluster 3, a population of cells shared between blood and tumor (Fig. 2c-d).

**Figure 4:**
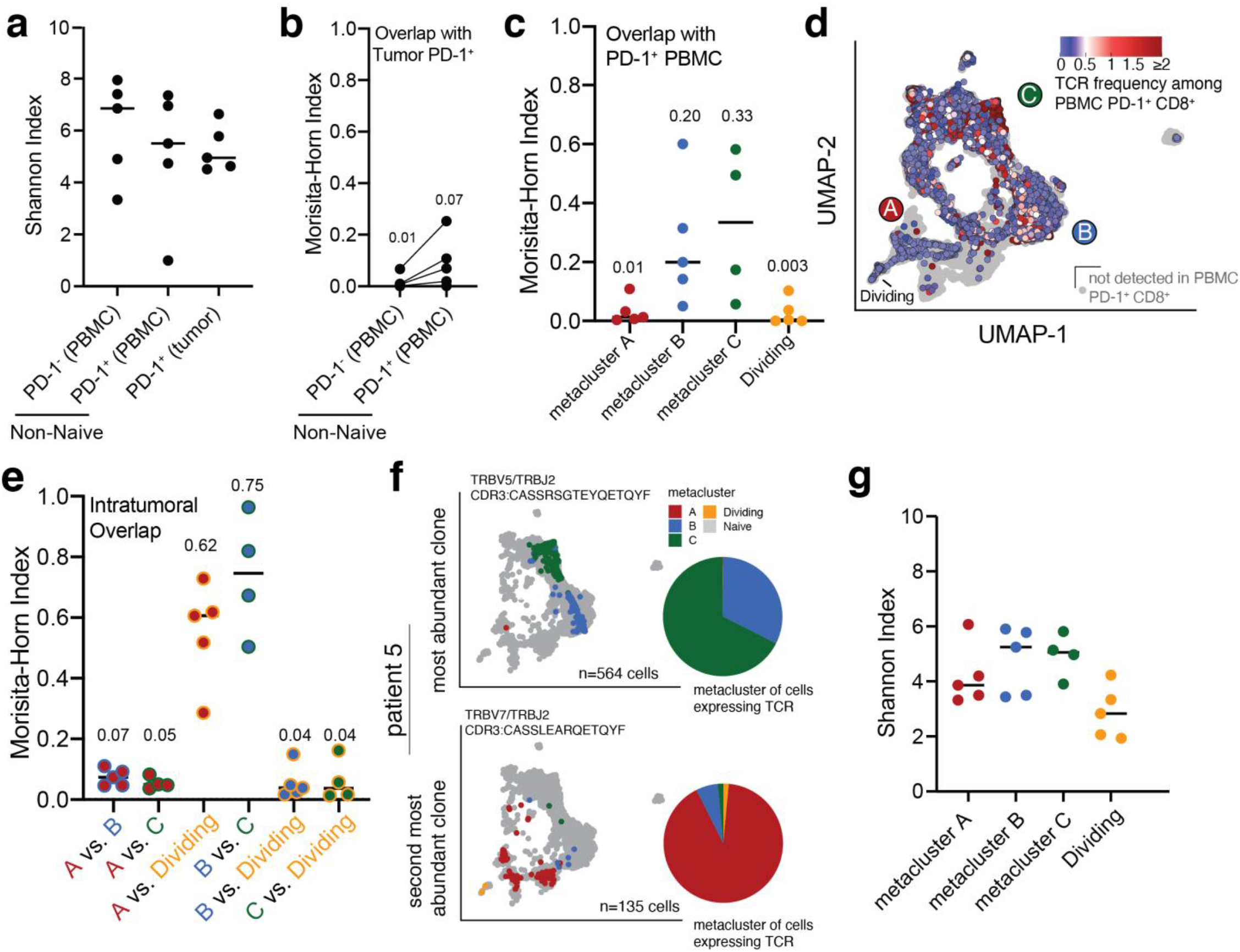
Dividing and terminally differentiated CD8^+^ T cells are clonally related to each other but not to other PD-1^+^ brain metastasis-infiltrating or circulating CD8^+^ T cells. (a) TCR diversity of circulating and brain metastasis-infiltrating CD8^+^ T cells. (b) Quantification of TCR overlap between tumor-infiltrating PD-1^+^ CD8^+^ T cells with circulating PD-1^−^ and PD-1^+^ cells from individual patients. (c) Quantification of TCR overlap between circulating PD-1^+^ CD8^+^ T cells and tumorin-filtrating PD-1^+^ CD8^+^ T cells within each metacluster. (d) UMAP colored by frequency of each cell’s TCR among circulating PD-1^+^ CD8^+^ T cells. (e) Quantification of TCR overlap between PD-1^+^ CD8^+^ T cells within each brain metastasis-infiltrating metacluster. (f) UMAP projection of cells expressing the most abundant (top) and second most abundant (bottom) TCR clone in patient 5, colored according to metacluster. Gray dots represent all other cells from the patient. Pie charts show distribution among metaclusters of cells expressing the TCR. All other patients are shown in Supplementary Figure 5. (g) TCR diversity of intratumoral metaclusters. In panels (a), (c), (e), and (g), lines indicate the median.

To interrogate the differentiation pathways of brain metastasis-infiltrating PD-1^+^ CD8^+^ T cells, we analyzed TCR overlap among scRNA-seq metaclusters. Terminally-differentiated and dividing cells (metaclusters A and D, respectively) had substantial TCR overlap with each other (Fig. 4e). However, these dividing and exhausted cells exhibited minimal TCR overlap with metaclusters B and C, which contained cells with a less exhausted phenotype (Fig. 4e-f, Supplementary Fig. 5). Most CD8^+^ T cell clones – and particularly the most abundant clones - within each patient’s tumor were mostly restricted to either an exhausted (metacluster A/D) or progenitor-like (metacluster B/C) phenotype (Fig. 4f, Supplementary Fig. 5), suggesting that these populations have largely unshared antigen-specificity. TCR diversity was also lower among terminally-differentiated cells (metacluster A/D) compared to cells in metaclusters B/C (Fig. 4g). Based on these data, we hypothesized that cells in metaclusters B and C may largely be memory bystander CD8^+^ T cells specific for non-tumor antigens and have become resident within the tumor following migration from the circulation.

### Brain metastasis-infiltrating bystander CD8^+^ T cells have phenotypic similarities to exhausted progenitor CD8^+^ T cells

Previous studies have identified tumor-infiltrating bystander CD8^+^ T cells^53,54^, but little is known about bystander infiltration of brain metastases. We queried the VDJdb, a database of TCRs with known specificity^55^, for matches with TCRs from our scRNA-seq data and identified one CMV-specific TCR in each of two patients (Supplementary Fig. 6). Both CMV-specific TCRs were expressed exclusively by cells in metaclusters B/C. To experimentally interrogate the abundance and phenotype of brain metastasis-infiltrating bystander CD8^+^ T cells, we *ex vivo* expanded PBMCs of four patients from whom scRNA-seq data were available. We stimulated these cells with a microbial peptide pool (CEFX), isolated IFNγ^+^ and IFNγ^−^ CD8^+^ T cells by FACS and performed TCR sequencing on each subset to identify TCRs that responded to CEFX stimulation with cytokine secretion and were thus microbe-specific (Fig. 5a and Supplementary Fig. 7).

**Figure 5:**
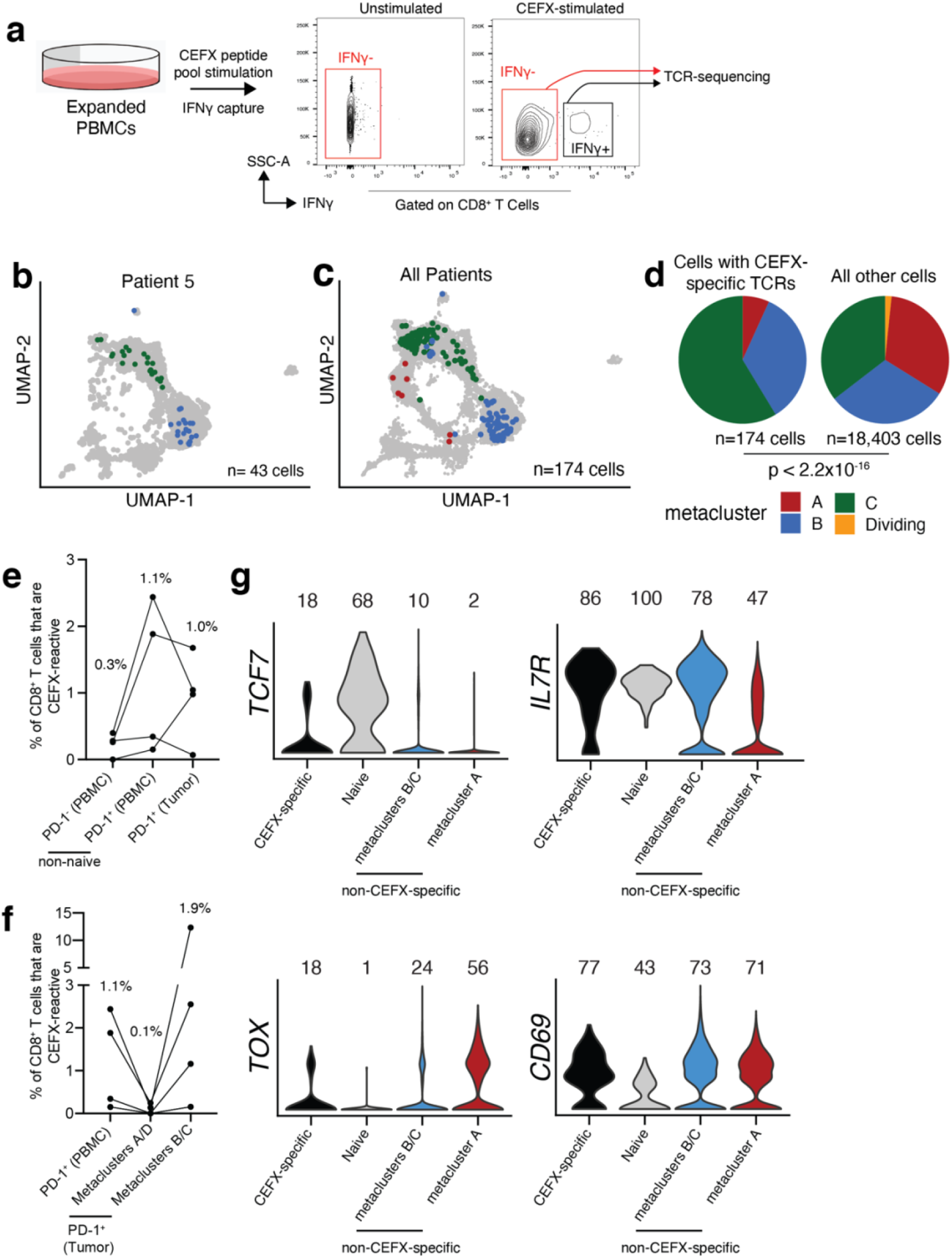
Microbe-specific CD8^+^ T cells are present in human brain metastases and are enriched in metaclusters B/C. (a) We performed IFN-γ capture on expanded, CEFX-stimulated PBMCs and subsequently sorted IFN-γ^+^ and IFN-γ^−^ CD8^+^ T cells for TCR sequencing to identify CEFX-specific TCRs. (b, c) Brain metastasis-infiltrating PD-1^+^ CD8^+^ T cells with CEFX-specific TCRs colored by phenotype on the UMAP for patient 5 (b) and all patients (c). (d) scRNA-seq phenotype of CEFX-specific (left) and all other (right) brain metastasis-infiltrating PD-1^+^ CD8^+^T cells. (e) Frequency of CEFX-specific cells among circulating and tumor-infiltrating CD8^+^ T cells. (f) Frequency of CEFX-specific cells among circulating PD-1^+^ CD8^+^ T cells and tumor-infiltrating PD-1^+^ CD8^+^ T cell subsets. (g) Expression of selected genes by CEFX-specific brain metastasis-infiltrating PD-1^+^ CD8^+^ T cells and all remaining cells by metacluster. In (e) and (f), medians are shown above each column. In (g), numbers indicate percentage of cells with measurable expression of the indicated marker. P value in (d) was calculated by Fisher’s Exact Test.

Comparison of TCR sequences from this assay and those identified from the scRNA-seq data above revealed that CEFX-specific cells ranged from 0.07% to 1.70% of brain metastasis-infiltrating PD-1^+^ CD8^+^ T cells. Phenotypically, CEFX-specific brain metastasis-infiltrating PD-1^+^ CD8^+^ T cells were found within metaclusters B/C in all four patients, where they were significantly enriched (Fig. 5b-d; Supplementary Fig. 8). A median of 1.9% of metacluster B/C cells were CEFX-specific, compared to 0.07% in metaclusters A and D, and 1.1% in total circulating PD-1^+^ CD8^+^ T cells (Fig. 5f). Of note, 12.3% of metacluster B/C cells from patient 17 were CEFX-specific. Importantly, some of these experimentally-validated bystander cells had a transcriptional phenotype similar to tumor-specific progenitor PD-1^+^ CD8^+^ T cells characterized in other studies,^41^ marked by expression of *IL7R* (CD127), *TOX*, and *TCF7* (TCF-1) (Fig. 5g; Supplementary Fig. 9). Of PD-1^+^ CD8^+^ T cells within the tumor, *IL7R* and *TCF7* expression was highest on CEFX-specific cells (Fig. 5).

Given the small number of known microbial T cell epitopes tested by this approach and the similar frequencies of bystander, non-tumor-specific cells between circulating and tumor-infiltrating PD-1^+^ T cells (Fig. 5e), the frequency of total bystander cells within human brain metastases is likely much higher. Additionally, given the enrichment of CEFX-specific cells in metaclusters B/C compared to both tumor-infiltrating terminally-differentiated cells and circulating cells (Fig. 5d), it is probable that many cells in metaclusters B/C are specific for non-tumor antigens and do not give rise to cells with a terminally-differentiated phenotype within the tumor. While these data do not preclude the presence of a small tumor-specific exhausted progenitor population within brain metastases, a large fraction of tumor-infiltrating TCF-1^+^ CD8^+^ T cells appear to be bystander cells specific for non-tumor antigen.

### CD8^+^ T cell phenotype is related to spatial distribution within the tumor

Given the divergent phenotypes and antigen-specificity of terminally-differentiated cells compared to other subsets within the tumor, we hypothesized that each subset of brain metastasis-infiltrating CD8^+^ T cells may be located within distinct regions of the TME and thus receiving different signals from surrounding tissue. To test this, we performed spatial transcriptomics, a method to measure gene expression *in situ*, on six brain metastasis tissue sections: one melanoma (Fig. 6), one renal cell carcinoma (Supplementary Fig. 10), one breast carcinoma (Supplementary Fig. 11), and three lung carcinomas (Supplementary Figs. 12-14). Capture spots were clustered based on gene expression and each cluster was annotated based on appearance in H&E staining.

**Figure 6:**
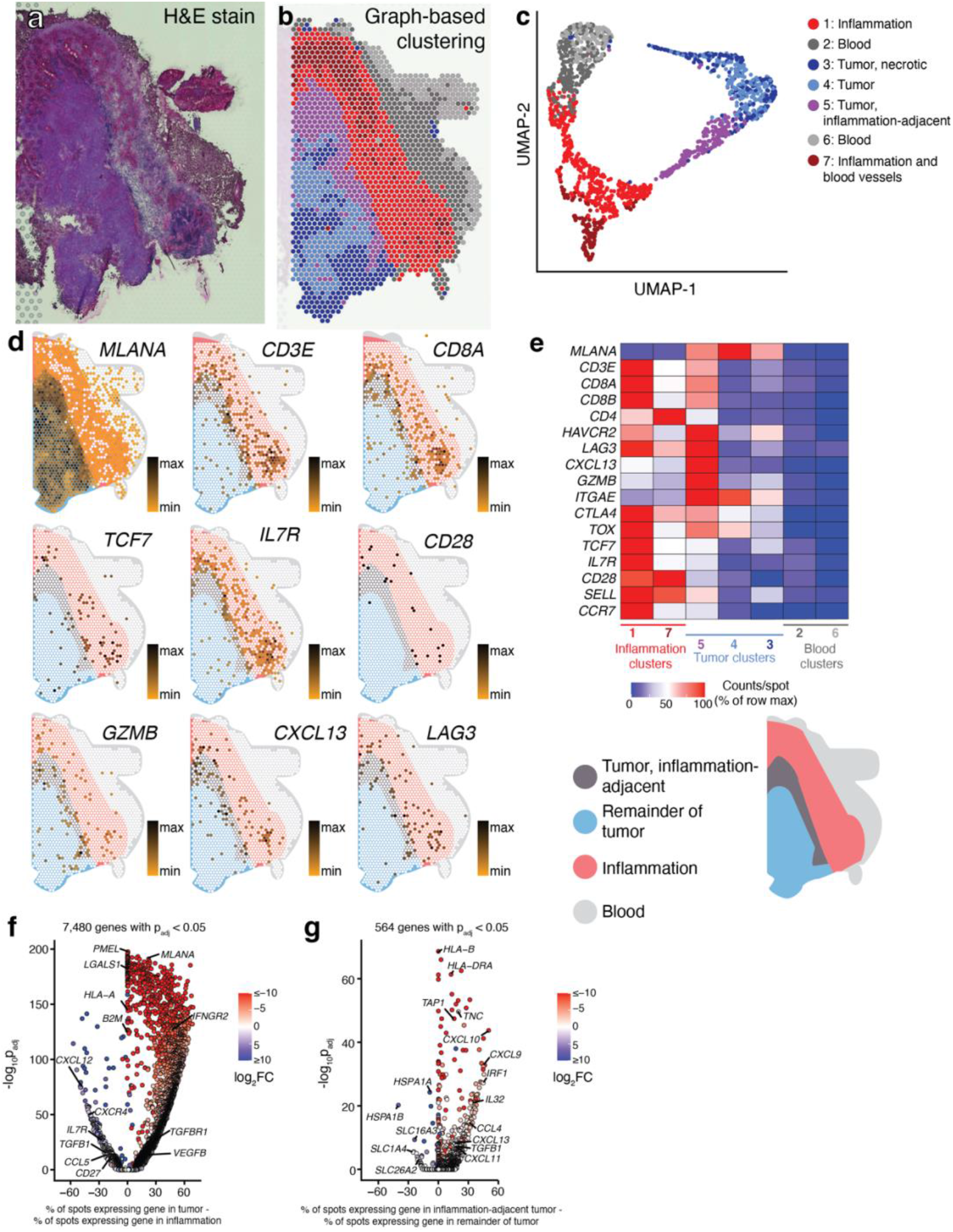
Genes associated with the terminally-differentiated CD8^+^ T cell phenotype are preferentially expressed within the tumor parenchyma of a melanoma brain metastasis. (a) Hematoxylin and eosin-stained section of a melanoma brain metastasis (patient 16). (b) Spatial location of capture spots, colored by transcriptional cluster. (c) UMAP and clustering of capture areas by transcriptional phenotype. (d) Expression of selected genes within the tissue section. White indicates no detection of the indicated gene in a given capture area. The darker the spot color, the greater the expression of the indicated gene. A spatial legend is given at bottom right. (e) Normalized expression density of selected genes in each tissue cluster. (f) Gene expression differences between tumor (clusters 3, 4, and 5) and peritumoral inflammation (clusters 1 and 7). (g) Differential gene expression between inflammation-adjacent tumor (cluster 5) and the remainder of tumor (clusters 3 and 4).

Tumor parenchyma — referring to regions of tumor cells — and surrounding stroma were readily differentiated by their gene expression profiles (Fig. 6a-b, Supplementary Figs. 10, 11, 13, and 14). For sections where tumor-normal boundaries were clearly visible, the number of differentially expressed genes between tumor parenchyma and surrounding stroma ranged from 3,914−7,480 (Fig. 6f, Supplementary Figs. 13c, 14e). In the two samples where tumor parenchyma and stroma were intermixed, the number of differentially expressed genes between the two regions was 637 and 2,327 (Supplementary Figs. 10e, 11e). One of the six tissue sections was comprised entirely of tumor parenchyma, precluding this analysis (Supplementary Fig. 12a-c).

The interplay between TME heterogeneity and immune infiltration was apparent. In the renal cell carcinoma case, “small nests” of tumor showed higher density of *CD8A, CD8B* and *CD4* expression than “large nests” of tumor, and transcript levels were even higher in regions of desmoplasia surrounding vessels (Supplementary Fig. 10c-d). Additionally, in two lung carcinoma samples where tumor parenchyma and brain tissue were visible, *CD3E* transcript levels were highest at the tumor interface with brain (Supplementary Figs. 13d and 14c-d). In the example of a melanoma brain metastasis tissue section where tumor parenchyma was surrounded by inflammation (patient 16), densities of *CD3E, CD4, CD8A*, and *CD8B* transcripts were highest in the peritumoral inflammation and in the directly adjacent tumor parenchyma (Fig. 6a-b, d-e). Together, these results are consistent with previous observations that immune cells are enriched in the peripheral region of brain metastases compared with the tumor core.^11^ In the melanoma sample, expression of genes associated with the terminally-differentiated phenotype of CD8^+^ T cells such as *HAVCR2* (TIM-3), *LAG3, CXCL13* and *GZMB*, was highest in the tumor parenchyma adjacent to inflammation, while expression of the progenitor markers *TCF7* (TCF-1) and *IL7R* (CD127) was highest in the inflammatory stroma suggesting that immune cell phenotype determines its location within the diverse TME (Fig. 6d-e).^11^ In the melanoma sample, expression of genes associated with the terminally-differentiated phenotype of CD8^+^ T cells such as *HAVCR2* (TIM-3), *LAG3, CXCL13* and *GZMB*, was highest in the tumor parenchyma adjacent to inflammation (Fig. 6d-e). Together, these data suggest that CD8^+^ T cell phenotype and location within the brain metastasis TME are linked.

### Terminally-differentiated CD8^+^ T cell clones are preferentially located in the tumor parenchyma

Genes that define these CD8^+^ T cell phenotypes may also be expressed by other cells within the TME, such as CD4^+^ T cells. Because we have shown that TCR clones in brain metastases are phenotypically restricted – that is, cells expressing a single TCR are predominately within scRNA-seq metaclusters A/D or B/C, but not both (Fig. 4e-f, Supplementary Fig. 5a, Fig. 7a) – localization of TCRs within the tumor would allow for visualization of specific CD8^+^ T cell phenotypes within the tumor. To determine whether there is spatial restriction of CD8^+^ T cells in the brain metastasis TME, we developed a method to amplify TCR transcripts from spatial transcriptomics gene expression libraries^56^. By linking TCR clones found in both spatial transcriptomics and scRNA-seq data, we could determine the location within the TME of CD8^+^ T cells with specific transcriptional phenotypes (Fig. 7b).

**Figure 7:**
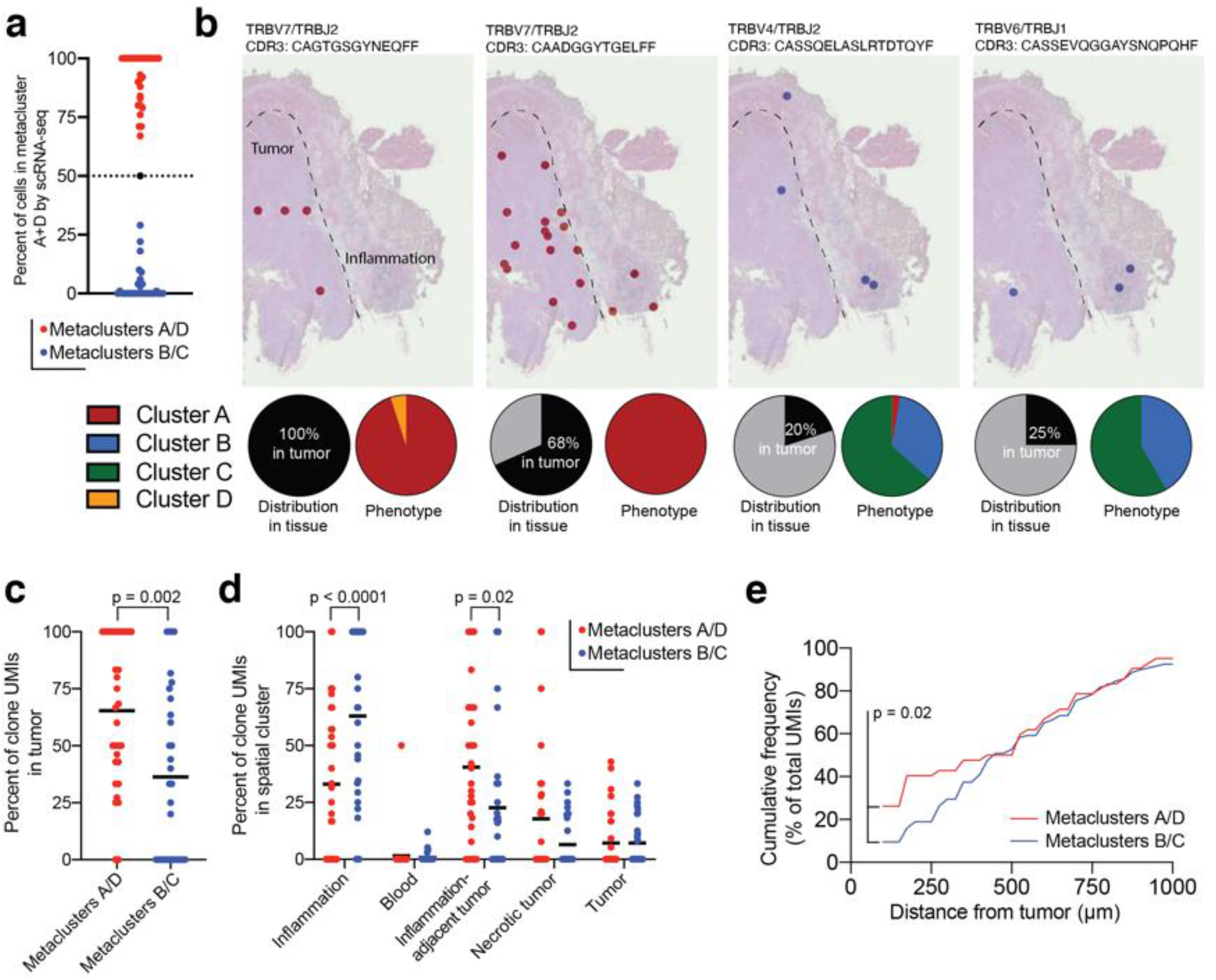
CD8^+^ T cell phenotype dictates location in the tumor microenvironment. (a) scRNA-seq phenotype of brain metastasis-infiltrating PD-1^+^ CD8^+^ T cells expressing TCRs identified by spatial transcriptomics. Each dot represents one TCR clone identified in both scRNA-seq and spatial transcriptomics data. For subsequent analysis, TCR clones were classified as metacluster A/Dividing clones (red) or metacluster B/C clones (blue) based on the scRNA-seq phenotype of cells expressing the clone. (b) Spatial location of selected TCR clones within tissue. Each dot represents a capture area in which at least one unique molecular identifier (UMI) for the indicated clone was found. Pie charts indicate percent of UMIs found in the tumor parenchyma (left) and percent of cells expressing the indicated TCR in each scRNA-seq metacluster (right). (c) Spatial distribution of UMIs from metacluster A/D and metacluster B/C clones in the tissue section shown in (b). Each dot is a single TCR clone. (d) Distribution of TCR UMIs in tissue clusters identified by spatial transcriptomics. (e) Cumulative frequency of UMIs outside the tumor parenchyma as a function of distance from the tumor border. The P-value in (c) was calculated by Mann-Whitney Test. P-values in (d) were calculated by two-way ANOVA with Sidak’s multiple comparisons test. The P-value in (e) was calculated by the Kolmogorov-Smirnov test.

Of the six tissues on which we performed spatial transcriptomics, scRNA-seq data from fresh tissue were available for two - patients 15 and 16, a lung carcinoma and melanoma sample, respectively. In the melanoma sample (patient 16), we observed that CD8^+^ T cell clones with a metacluster A/Dividing phenotype were predominantly located in the tumor parenchyma, while CD8^+^ T cell clones with a metacluster B/C phenotype were predominantly found in the peritumoral inflammation (Fig. 7b-d). Not only were metacluster A/Dividing TCRs enriched in the tumor parenchyma, but TCR clones with this phenotype found outside of the tumor were also preferentially located closer to the tumor boundary compared to those with a metacluster B/C phenotype (Fig. 7e). These findings were confirmed in the lung carcinoma brain metastasis (patient 15) where the entire tissue section was tumor parenchyma and inhabited only by TCRs expressed by metacluster A/Dividing cells (Supplementary Fig. 12e-j).

Given this preference of specific CD8^+^ T cell subsets for particular locations within the TME, we sought to determine whether they receive distinct signaling inputs based on their location. We therefore compared the transcriptional profiles between different spatial gene expression clusters of the tumor. In the renal cell carcinoma sample (Patient 24), gene expression varied with tumor architecture: 305 genes were differentially expressed between small and large nests of tumor (Supplementary Fig. 10f-h). MHC I expression was higher within small tumor nests compared to large tumor nests (Supplementary Fig. 10f), suggesting that CD8^+^ T cells within the same tumor may receive different levels of TCR stimulation based on their location within the parenchyma. In the melanoma sample (Patient 16), 564 genes were differentially expressed between the region of tumor adjacent to inflammation and the remainder of the tumor parenchyma (Fig. 6g). Transcript levels of MHC I and MHC II were higher in the peripheral, inflammation-adjacent tumor compared to the remainder of the parenchyma, suggesting that TCR stimulation of tumor-specific cells is greatest in this region (Fig. 6g). Conversely, in the breast carcinoma sample (patient 24), MHC I was highly downregulated in capture spots containing tumor parenchyma, potentially indicating limited tumor-associated antigen presentation to CD8^+^ T cells in this patient (Supplementary Fig. 11g). In the melanoma sample (patient 16), *CXCL9, CXCL10, CXCL11* and *CXCL13* as well as TGF-β were higher at the periphery, indicating that T cells in this region are subject to a unique chemokine and cytokine milieu compared to those deeper within the tumor (Fig. 6g).

Differences in cytokine and chemokine expression between bulk tumor parenchyma and surrounding tissue were also striking (Supplementary Fig. 15a). As examples, *TGFB1* (encoding TGF-β) was enriched in the stroma in three samples, whereas its receptor was more highly expressed predominately in the parenchyma (Supplementary Fig. 15a-b). The IFN-γ receptor subunit *IFNGR2* was elevated in the parenchyma of four of five tumors (Supplementary Fig. 15a,c). *VEGFA* and *VEGFB* were upregulated in the parenchyma all five tumors (Supplementary Fig. 15a,d). Given the differential localization of phenotypically and clonally restricted CD8^+^ T cell subsets in the tumor, our results together indicate that antigen signaling (or lack thereof) drives these distinct CD8^+^ T cell subsets to specific niches within brain metastases, where they receive markedly different signaling inputs.

## Discussion

In this work, we describe the CD8^+^ T cell infiltrate of human brain metastases, a significant cause of morbidity and mortality in many cancer types. The blood-brain-barrier (BBB), which maintains the unique immune privileged environment of the brain, appears to break down in brain metastases. The resulting blood-tumor-barrier (BTB) is more permeable but maintains some features of the BBB and varies with tumor type^6^. Although brain metastases show some response to ICB, the semi-privileged immune environment created by the BTB could, in theory, restrict entry and/or maintenance of immune cells into the brain metastasis TME. Here, we show that human brain metastases are well-infiltrated by CD8^+^ T cells, although the degree of infiltration varies among patients.

We find that distinct CD8^+^ T cell subsets populate brain metastases compared to patient-matched blood. Most CD8^+^ T cells within the tumor are PD-1^+^, and our scRNA-seq analyses clustered these PD-1^+^ cells into dividing and terminally-differentiated cells – which are clonally related – and two less exhausted subsets of cells which share TCR overlap with each other but not with dividing or terminally-differentiated cells. Overall, TCR overlap is low between tumor-infiltrating and circulating PD-1^+^ CD8^+^ T cells, consistent with the apparent absence of circulating tumor-specific CD8+ T cells in HPV+ head and neck cancer patients^35^. However, less exhausted tumor-infiltrating cells do have meaningful clonal overlap with circulating cells, whereas terminally-differentiated tumor-infiltrating cells have very little. Notably, we find that brain metastases contain non-tumor-specific CD8^+^ T cells, these cells express markers also found on exhausted progenitor populations, and are found in similar frequencies in tumor-infiltrating and circulating PD-1^+^ CD8^+^ T cells. Finally, we link these phenotypically- and clonally-restricted CD8^+^ T cells to discrete spatial preferences within the TME. CD8^+^ T cell clones linked to exhaustion are enriched within the tumor parenchyma, where the local cytokine and chemokine milieu varies dramatically from the surrounding stroma where less exhausted CD8^+^ T cell clones are found. Based on these findings, our data support a model in which CD8^+^ T cells infiltrate the TME of brain metastases in an antigen-independent manner^38^. Once CD8^+^ T cells are retained in the tumor, antigen signaling, or lack thereof, drives recruitment of CD8^+^ T cells to spatial niches within the TME. Within each TME niche, CD8^+^ T cell phenotype evolves with inputs from the surrounding cytokine milieu and, depending on antigen specificity, continued TCR stimulation.

One key question this work addresses is the role of TCF-1^+^ CD8^+^ T cells in human brain metastases. TCF-1 is expressed on antigen-specific exhausted progenitor CD8^+^ T cells in mouse models of cancer and chronic infection^23,41^, and TCF-1 has been used as a marker of exhausted progenitor CD8^+^ T cells in tumor immunology studies^32–34^. However, TCF-1 is also expressed on other subsets of CD8^+^ T cells such as naïve and memory cells^57^. We find that a minority of TCF-1^+^ CD8^+^ T cells within brain metastases co-express TOX and PD-1, two proteins also expressed on exhausted, tumor-specific CD8^+^ T cells^35,43–45,49–51^. Crucially, we show that there are bystander cells specific for microbial antigens infiltrating brain metastases, and that a subset of these cells share phenotypic characteristics - such as TCF-1 expression - with exhausted progenitor CD8^+^ T cells. Bystander clones are present at similar frequencies in circulating and tumor-infiltrating PD-1^+^ CD8^+^ T cells. Based on these data, we propose that many of TCF-1^+^ PD-1^+^ CD8^+^ T cells in the brain metastasis TME are bystanders. These cells may be recruited to the tumor due to increased expression of cytokines and pro-inflammatory signaling molecules rather than through an antigen-driven process. As such, increased density of TCF-1^+^ CD8^+^T cells in the brain metastasis TME may indicate a more inflamed tumor rather than a large population of tumor-specific exhausted progenitor CD8^+^ T cells.

We show that less-exhausted brain metastasis-infiltrating CD8^+^ T cells are preferentially retained in the stroma surrounding the tumor parenchyma. In contrast, the terminally-differentiated population is preferentially located within the tumor parenchyma itself. Given the limited clonal overlap between bystander cells – which adopt a less-exhausted phenotype - and the terminally-differentiated population, it is likely antigen stimulation that drives terminally-differentiated PD-1^+^ CD8^+^ T cells to the tumor parenchyma. Within the tumor parenchyma, these cells receive distinct signaling inputs compared to CD8^+^ T cells within to the stroma, likely promoting their acquisition of the terminally-differentiated phenotype. We show that VEGF expression, for example, is enriched in the tumor parenchyma compared to the stroma; VEGF signaling has been found to promote PD-1, CTLA-4, TIM-3, and TOX expression on CD8^+^ T cells^58^. This is consistent with the higher expression of these molecules we observed on terminally-differentiated CD8^+^ T cells.

The dividing CD8^+^ T cell metacluster identified in our scRNA-seq data could represent an intermediate differentiation state between tumor-specific TCF-1^+^ exhausted progenitor and terminally-differentiated cells. These dividing cells in human brain metastases share a gene signature with transitory antigen-specific CD8^+^ T cells that are an intermediate differentiation state between lymphoid-resident exhausted progenitor cells and nonlymphoid-resident, terminally-differentiated cells in the LCMV mouse model of T cell exhaustion^24,26^. Work in mouse tumor models has shown that tumor-specific exhausted progenitor CD8^+^ T cells are present in tumor-draining lymph nodes, and that they are clonally related to tumor-infiltrating, terminally-differentiated CD8^+^ T cells^39–41^. Further, the maintenance of antigen-specific CD8^+^ T cells within mouse models requires the migration of lymph-node resident progenitor cells to the tumor; intratumoral TCF-1^+^ tumor-specific cells are not a self-sustaining population^40^. In the case of metastatic cancer, there may be numerous anatomic sites of tumor draining lymph nodes depending on the burden of disease, all of which could contain tumor-specific exhausted progenitor CD8^+^ T cells. A lymphatic system draining the brain has recently been described^59–61^ and suggests that cervical lymph nodes may also serve as a reservoir of brain metastasis-specific progenitor CD8^+^ T cells. Our data do not preclude the presence of a small population of tumor-specific progenitor exhausted CD8^+^ T cells within the brain metastasis TME but suggest that these other sites may be an important reservoir of tumor-specific exhausted progenitors.

Our findings have a number of therapeutic implications. First, the dense infiltration of brain metastases by CD8^+^ T cells provides a rational basis for the use and development of immunotherapies in the brain metastasis setting. Next, our data support the use of combination therapies with PD-1 pathway blockade to enhance rescue of exhausted, tumor-specific CD8^+^ T cells by targeting additional inhibitory molecules expressed on brain metastasis-infiltrating terminally-differentiated CD8^+^ T cells, such as CTLA-4 and LAG3^62,63^. These terminally-exhausted CD8^+^ T cells within the TME may re-gain effector function with PD-1 blockade and/or combination therapies that lead to enhanced tumor cell killing^31^. Finally, targeting the unique signaling niches in which brain metastasis-infiltrating CD8^+^ T cells reside provides an additional opportunity to harness the immune system to improve disease control in the brain. Together our findings support the continued development of immunotherapeutic strategies that harness the anti-tumor efficacy of brain metastasis-infiltrating CD8^+^ T cells.

## Acknowledgements

We are grateful to the Emory University School of Medicine Flow Cytometry Core for assistance with cell sorting and to Kathryn Pellegrini of the Yerkes Nonhuman Primate (NHP) Genomics Core for assistance with scRNA-seq captures. L.J.S. is supported by a Conquer Cancer Young Investigator Award, a Radiological Society of North America Resident Research Award, and the Nell W. and William Simpson Elkin Fellowship from Emory University. R.A. is supported by R01AI030048 and P01AI056299 from the National Institutes of Health (NIH). W.H.H. is supported by a Cancer Research Institute Irvington Fellowship from the Cancer Research Institute and K99AI153736 from the NIH. The Yerkes NHP Genomics Core is supported in part by NIH P51OD011132, and sequencing data was acquired on an Illumina NovaSeq6000 funded by NIH S10OD026799. Research reported in this publication was also supported by the Ambrose Monell Foundation, the Oliver S. and Jennie R.

Donaldson Charitable Trust, the National Cancer Institute (NCI) of the NIH under Award Number P50CA217691 and by the Pediatrics/Winship Flow Cytometry Core of Winship Cancer Institute of Emory University, Children’s Healthcare of Atlanta and NIH/NCI under award number P30CA138292. The content is solely the responsibility of the authors and does not necessarily represent the official views of the National Institutes of Health or the American Society of Clinical Oncology® or Conquer Cancer®.

## Data deposition

Raw and processed data for scRNA-seq have been deposited in the Gene Expression Omnibus (GEO) under accession GSE179373. Raw and processed data for the gene expression spatial transcriptomics experiments have been deposited in the GEO with accession GSE179572. Reads from targeted TCR-sequencing for spatial transcriptomics have been deposited in the Sequence Read Archive under BioProject PRJNA742564.

## Methods

### Patients and sample acquisition (blood and tumor)

All brain metastases from immunotherapy-naïve patients resected at Emory University Hospital during the collection period were included unless consent was not obtained, there was insufficient tissue, or in some cases the surgery was performed emergently after hours. Blood samples were collected during surgery in BD Vacutainer lithium heparin tubes. Tumor and blood samples were held at 4 °C until retrieved for processing, typically within 1 hour after resection. Nearly all patients with surgically-resected brain metastases are treated with the immunosuppressive glucocorticoid dexamethasone prior to surgery. Standard dexamethasone administration was a 10 mg loading dose followed by 4 mg every 6 hours until surgery. The duration of dexamethasone therapy prior to surgery was not correlated with CD45^+^ lymphocyte or CD8^+^ T cell infiltration of brain metastases in our cohort (Supplementary Fig. 16).

Experiments were carried out with the approval of the Emory University Institutional Review Board under protocols IRB00045732, IRB00095411, and STUDY00001995.

### Tissue processing and cell extraction

Tumors were weighed, cut into small pieces, and then incubated in Leibovitz media with collagenase I, II, and IV, elastase and DNAse for 60 minutes, shaking at 37 °C. Tissue and media were then passed through a 70 μm single-cell strainer and cells were pelleted by centrifugation. The pellet was resuspended in 44% Percoll, underlaid with 67% Percoll, and centrifuged. Immune cells were collected from the gradient interface and washed with 2% FBS in PBS. Washed cells were resuspended in 1-2 mL of PBS containing 2% FBS. 10 μL of cells were stained for 20 minutes with anti-CD45 and anti-CD8 antibodies. CountBright counting beads (Thermo Fisher) were added to stained cells and the sample was analyzed on a BD LSR II flow cytometer to determine the absolute number of CD45^+^ lymphocytes and CD8^+^ T cells isolated. For isolation of circulating immune cells, a lymphocyte separation medium (Corning catalog #25-072-CV) gradient was performed according to the manufacturer’s instructions. With the exception of scRNA-seq samples (see below), cells were then frozen at −80 °C in 10% DMSO in FBS and subsequently transferred to liquid nitrogen for long-term storage. For scRNA-seq, cells were immediately stained with antibodies for the sort as described below.

### Flow cytometry

For scRNA-seq and TCR sequencing, freshly-isolated tumor-infiltrating and circulating immune cells were stained with extracellular antibodies for 30 minutes, washed with 2% FBS in PBS and sorted on a BD FACS ARIA II in the Emory Flow Cytometry Core Facility. Tumor-infiltrating PD-1^+^ CD8^+^ T cells were submitted to the Emory Yerkes Genomics Core where gene expression and TCR sequence libraries were generated with a 10X Genomics Chromium controller. DNA was extracted from circulating PD-1^+^ and PD-1^−^ CD8^+^ T cells using the All-Prep DNA/RNA Micro Kit from Qiagen and sent to Adaptive Biotechnologies for survey-level TCRβ sequencing.

For high-parameter flow cytometry, frozen cells were quickly thawed in a 37 °C water bath, washed with pre-warmed (37 °C) 10% FBS in RPMI, and resuspended in staining buffer (PBS with 2% FBS). Staining was performed at room temperature. Washed cells were stained first with Zombie NIR viability dye for 30 minutes, then extracellular antibodies were added in BD Horizon Brilliant Stain Buffer for 30 minutes. Cells were washed twice in staining buffer. The eBioscience Foxp3 Transcription Factor Staining Buffer Set was then used for fixation and permeabilization according to the manufacturer’s protocols. Cells were then stained with intracellular antibodies for 30 minutes, washed twice with permeabilization buffer, stained with secondary antibodies for 30 minutes, washed with permeabilization buffer once and staining buffer once, then resuspended in staining buffer for data acquisition. Data was acquired on a Cytek Aurora flow cytometer in the Winship Pediatrics Flow Cytometry Core. Data were analyzed in FlowJo, using the FlowSOM and UMAP plugins^64,65^. Summary graphs and statistics were performed in GraphPad Prism v8.

### PBMC expansion, CEFX stimulation, and IFNγ capture

PBMCs were quickly thawed, washed, and counted in pre-warmed 10% FBS in RPMI. Cells were expanded as described previously^35^. Briefly, cells were resuspended in complete CTS media containing CTS OpTmizer (Gibco) with CTS supplement, L-glutamine, Penicillin/Streptomycin, Human AB serum (Sigma), recombinant IL-2, IL-7, and IL-15 (Peprotech), and CEFX Ultra SuperStim peptide pool (JPT). Cells were plated and incubated at 37 °C for five days and then split with the above CTS complete media and cytokines, without the addition of the CEFX peptides. Five days later, cells were washed and rested overnight at 37 °C in complete CTS media without cytokines. The next day, the CEFX Ultra SuperStim peptide pool was added to cells at a final concentration of 2.5 μg/mL. An equal volume of DMSO was added to unstimulated cells. Cells were incubated 5 hours at 37 °C. Manufacturer’s protocols were then followed for the Miltenyi cytokine secretion assay (catalog #130-054-202) using the cytokine catch reagent and cytokine detection antibody (IFN-γ PE) to label cells secreting IFN-γ. Cells were then stained for viability, CD3, and CD8 and sorted on a BD FACS ARIA II in the Emory Flow Cytometry Core (Supplementary Figure 7). DNA was isolated from sorted IFNγ^+^ and IFNγ^−^ CD8^+^ T cells using the All-Prep DNA/RNA Micro Kit from Qiagen and sent to Adaptive Biotechnologies for survey-level TCRβ sequencing. A clone was considered CEFX-specific if found at least twice in the IFN γ^+^ population and with a frequency ≥5x higher in IFNγ^+^ cells compared to IFNγ^−^ cells.

### Single cell RNA-sequencing, single cell TCR analysis, and peripheral TCR profiling

Single-cell gene expression and VDJ (paired TCRα/β) libraries were generated by the Yerkes Nonhuman Primate Genomics Core from freshly-isolated tumor-infiltrating PD-1^+^ CD8^+^ T cells and matched naïve circulating CD8+ T cells isolated by FACS and mixed at a 10:1 ratio. TILs and naive circulating cells were stained with TotalSeq hashing antibodies (BioLegend) prior to combining for cell capture and library preparation. TCR sequencing (TCRβ only) from sorted circulating cells was performed by Adaptive Biotechnologies.

Single-cell gene expression data were aligned and TCR sequences determined with CellRanger version 4 (10X Genomics). Outlier cells with high numbers of reads originating from mitochondrial genes and presumed doublets were excluded from the dataset, and genes encoded on the Y or mitochondrial chromosomes were excluded from gene expression analysis. Data were normalized and scaled with the Seurat package in R^66^ and plots made with ggplot2^67^ or GraphPad Prism. Shared nearest neighbor clustering was performed in Seurat with 100 neighbors, 21 principal components, and a resolution of 0.9. UMAP dimensionality reduction was performed with 100 neighbors, 21 principal components, and a minimum distance of 0. Seurat’s BuildClusterTree with identical parameters was used for was used to create a phylogenetic tree of identified clusters. Gene set enrichment analysis (GSEA) was performed with the fgsea package in R^68^, using sign(FC)*-log_10_p_adj_ from Seurat’s FindMarkers function as the ranking statistic. Morisita-Horn indices were calculated with the divo package in R^69^. Shannon indices were calculated in R with the vegan package^70^.

For TCR analysis, cells or sequencing reads with identical *TRBV* and *TRBJ* gene families and identical TCRβ CDR3 amino acid sequences were considered to originate from same clone. Cells from scRNA-seq with undetermined TCRβ clonotypes but known TCRα sequences were assigned to a clonotype if all other cells with the TCRα were paired with a single TCRβ clone. To search for cells with known antigen specificity, we queried the VDJdb^71^ (accessed February 2021) with paired TCRα/β sequences from our scRNA-seq data, requiring identical TCRα and TCRβ CDR3 sequences as well as exact matches for *TRAV, TRAJ, TRBV, TRBJ* genes to be considered a T cell with known antigen specificity. This resulted in the identification of two clones specific for the CMV protein IE1.

### Spatial transcriptomics

Surgically resected tissue was embedded in OCT and immediately flash frozen in a dry ice/2-methylbutane bath. Sections 10 μm thick were placed onto a Visium Gene Expression slide (10X Genomics PN-1000184) and stored at −80 °C for up to one week. Slides were subsequently H&E stained according to the manufacturer’s instructions and imaged with a Lionheart Microscope (Biotek) at 10X magnification. Tissue permeabilization, reverse transcription, second strand synthesis, and cDNA amplification was performed according to the manufacturer’s instructions. 25% of amplified cDNA was used for gene expression library preparation; libraries were sequenced on a NovaSeq 6000 instrument at the Yerkes Nonhuman Primate Genomics Core.

For TCR library preparation, 5 μl of amplified cDNA was used as template in a 35-cycle PCR reaction using 45 previously-described^72^ pooled *TRBV* forward primers and the Illumina read 1 reverse primer. Partial Illumina read 2 sequences (5’-GTGACTGGAGTTCAGACGTGTGCTCTTCCGATCT-3’) were added to the 5’ end of each *TRBV* forward primer. PCR product was purified without fragmentation using SPRIselect beads and quantified using a Qubit 1X dsDNA HS Assay Kit (Thermo Fisher).

Sample index PCR was performed with the 10X Genomics Library Construction Kit (PN-1000190) using primers from the 10X Genomics Dual Index Kit TT Set A (PN-1000215) according to the manufacturer’s instructions (protocol CG000239, 10X Genomics). Libraries were again bead purified and sequenced on an Illumina MiSeq instrument at the Yerkes Nonhuman Primate Genomics Core. The MiXCR^73^ analyze pipeline was performed on read 2 sequences, and supporting reads for each clonotype were written with the exportReadsForClones command. The UMI and spatial barcodes were extracted from the paired read for each supporting sequencing read. A detailed protocol for obtaining TCR sequences from spatial transcriptomics data is available in the accompanying manuscript^56^.

Loupe Browser (10X Genomics) was to visualize data for pathology review; tissue regions were called by graph-based clustering and annotated within Loupe Browser. Detailed analysis and visualization were conducted in R with the Seurat package^66^. Genes were considered below the limit of detection if expressed below 10 counts or were expressed in two or fewer spots. For analyses of boundary and tumor gene expression (Supplementary Fig. 15) the following clusters were used: sample 16, clusters 5 and 3/4 (boundary and tumor, respectively); sample 19, clusters 3 and 5; sample 24, clusters 4 and 1/2/3; sample 26, clusters 6 and 1/2/3; sample 27, clusters 7 and 2/3/4/5/6.

## References

1 Achrol, A. S. et al. Brain metastases. Nature Reviews Disease Primers 5, doi:10.1038/s41572-018-0055-y (2019).

2 Singh, R. et al. Epidemiology of synchronous brain metastases. Neuro-Oncology Advances 2, doi:10.1093/noajnl/vdaa041 (2020).

3 Aquilanti, E. & Brastianos, P. K. Immune Checkpoint Inhibitors for Brain Metastases: A Primer for Neurosurgeons. Neurosurgery 87, E281–E288, doi:10.1093/neuros/nyaa095 (2020).

4 Lin, X. & DeAngelis, L. M. Treatment of Brain Metastases. Journal of clinical oncology: official journal of the American Society of Clinical Oncology 33, 3475–3484, doi:10.1200/jco.2015.60.9503 (2015).

5 Forrester, J. V., McMenamin, P. G. & Dando, S. J. CNS infection and immune privilege. Nature Reviews Neuroscience 19, 655–671, doi:10.1038/s41583-018-0070-8 (2018).

6 Arvanitis, C. D., Ferraro, G. B. & Jain, R. K. The blood–brain barrier and blood–tumour barrier in brain tumours and metastases. Nature Reviews Cancer 20, 26–41, doi:10.1038/s41568-019-0205-x (2020).

7 Nduom, E. K., Yang, C., Merrill, M. J., Zhuang, Z. & Lonser, R. R. Characterization of the blood-brain barrier of metastatic and primary malignant neoplasms. Journal of Neurosurgery 119, 427–433, doi:10.3171/2013.3.jns122226 (2013).

8 Berghoff, A. S. et al. Density of tumor-infiltrating lymphocytes correlates with extent of brain edema and overall survival time in patients with brain metastases. Oncoimmunology 5, e1057388, doi:10.1080/2162402x.2015.1057388 (2016).

9 Duchnowska, R. et al. Immune response in breast cancer brain metastases and their microenvironment: the role of the PD-1/PD-L axis. Breast Cancer Research 18, doi:10.1186/s13058-016-0702-8 (2016).

10 Berghoff, A. S. et al. Tumour-infiltrating lymphocytes and expression of programmed death ligand 1 (PD-L1) in melanoma brain metastases. Histopathology 66, 289–299, doi:10.1111/his.12537 (2015).

11 Amit, M. et al. Characterization of the melanoma brain metastatic niche in mice and humans. Cancer Medicine 2, 155–163, doi:10.1002/cam4.45 (2013).

12 Friebel, E. et al. Single-Cell Mapping of Human Brain Cancer Reveals Tumor-Specific Instruction of Tissue-Invading Leukocytes. Cell, doi:https://doi.org/10.1016/j.cell.2020.04.055 (2020).

13 Mansfield, A. S. et al. Contraction of T cell richness in lung cancer brain metastases. Scientific Reports 8, doi:10.1038/s41598-018-20622-8 (2018).

14 Lau, L. L., Jamieson, B. D., Somasundaram, T. & Ahmed, R. Cytotoxic T-cell memory without antigen. Nature 369, 648–652, doi:10.1038/369648a0 (1994).

15 Wherry, E. J. T cell exhaustion. Nature Immunology 12, 492–499, doi:10.1038/ni.2035 (2011).

16 Wherry, E. J. et al. Molecular signature of CD8+ T cell exhaustion during chronic viral infection. Immunity 27, 670–684, doi:10.1016/j.immuni.2007.09.006 (2007).

17 Barber, D. L. et al. Restoring function in exhausted CD8 T cells during chronic viral infection. Nature 439, 682–687, doi:10.1038/nature04444 (2006).

18 Topalian, S. L. et al. Safety, Activity, and Immune Correlates of Anti-PD-1 Antibody in Cancer. New England Journal of Medicine 366, 2443–2454, doi:10.1056/nejmoa1200690 (2012).

19 Zaretsky, J. M. et al. Mutations Associated with Acquired Resistance to PD-1 Blockade in Melanoma. New England Journal of Medicine 375, 819–829, doi:10.1056/nejmoa1604958 (2016).

20 Ott, P. A., Hodi, F. S., Kaufman, H. L., Wigginton, J. M. & Wolchok, J. D. Combination immunotherapy: a road map. Journal for immunotherapy of cancer 5, doi:10.1186/s40425-017-0218-5 (2017).

21 He, R. et al. Follicular CXCR5-expressing CD8+ T cells curtail chronic viral infection. Nature 537, 412–416, doi:10.1038/nature19317 (2016).

22 Utzschneider, D. T. et al. T Cell Factor 1-Expressing Memory-like CD8+ T Cells Sustain the Immune Response to Chronic Viral Infections. Immunity 45, 415–427, doi:10.1016/j.immuni.2016.07.021 (2016).

23 Im, S. J. et al. Defining CD8+ T cells that provide the proliferative burst after PD-1 therapy. Nature 537, 417–421, doi:10.1038/nature19330 (2016).

24 Hudson, W. H. et al. Proliferating Transitory T Cells with an Effector-like Transcriptional Signature Emerge from PD-1(+) Stem-like CD8(+) T Cells during Chronic Infection. Immunity 51, 1043–1058.e1044, doi:10.1016/j.immuni.2019.11.002 (2019).

25 Zander, R. et al. CD4+ T Cell Help Is Required for the Formation of a Cytolytic CD8+ T Cell Subset that Protects against Chronic Infection and Cancer. Immunity 51, 1028–1042.e1024, doi:10.1016/j.immuni.2019.10.009 (2019).

26 Im, S. J., Konieczny, B. T., Hudson, W. H., Masopust, D. & Ahmed, R. PD-1+ stemlike CD8 T cells are resident in lymphoid tissues during persistent LCMV infection. Proceedings of the National Academy of Sciences 117, 4292–4299, doi:10.1073/pnas.1917298117 (2020).

27 Yan, Y. et al. CX3CR1 identifies PD-1 therapy–responsive CD8+ T cells that withstand chemotherapy during cancer chemoimmunotherapy. JCI Insight 3, doi:10.1172/jci.insight.97828 (2018).

28 Yamauchi, T. et al. T-cell CX3CR1 expression as a dynamic blood-based biomarker of response to immune checkpoint inhibitors. Nature Communications 12, doi:10.1038/s41467-021-21619-0 (2021).

29 Pauken, K. E. et al. Single-cell analyses identify circulating anti-tumor CD8 T cells and markers for their enrichment. Journal of Experimental Medicine 218, doi:10.1084/jem.20200920 (2021).

30 Hashimoto, M. et al. CD8 T Cell Exhaustion in Chronic Infection and Cancer: Opportunities for Interventions. Annual review of medicine 69, 301–318, doi:10.1146/annurev-med-012017-043208 (2018).

31 Voabil, P. et al. An ex vivo tumor fragment platform to dissect response to PD-1 blockade in cancer. Nature Medicine 27, 1250–1261, doi:10.1038/s41591-021-01398-3 (2021).

32 Jansen, C. S. et al. An intra-tumoral niche maintains and differentiates stem-like CD8 T cells. Nature 576, 465–470, doi:10.1038/s41586-019-1836-5 (2019).

33 Brummelman, J. et al. High-dimensional single cell analysis identifies stem-like cytotoxic CD8+ T cells infiltrating human tumors. Journal of Experimental Medicine 215, 2520–2535, doi:10.1084/jem.20180684 (2018).

34 Sade-Feldman, M. et al. Defining T Cell States Associated with Response to Checkpoint Immunotherapy in Melanoma. Cell 175, 998–1013.e1020, doi:10.1016/j.cell.2018.10.038 (2018).

35 Eberhardt, C. S. et al. Functional HPV-specific PD-1+ stem-like CD8 T cells in head and neck cancer. Nature (In press).

36 Wang, H.-f. et al. The Double-Edged Sword—How Human Papillomaviruses Interact With Immunity in Head and Neck Cancer. Frontiers in immunology 10, doi:10.3389/fimmu.2019.00653 (2019).

37 Caushi, J. X. et al. Transcriptional programs of neoantigen-specific TIL in anti-PD-1-treated lung cancers. Nature, doi:10.1038/s41586-021-03752-4 (2021).

38 Oliveira, G. et al. Phenotype, specificity and avidity of antitumour CD8+ T cells in melanoma. Nature, doi:10.1038/s41586-021-03704-y (2021).

39 Buchwald, Z. S. et al. Tumor-draining lymph node is important for a robust abscopal effect stimulated by radiotherapy. Journal for immunotherapy of cancer 8, doi:10.1136/jitc-2020-000867 (2020).

40 Connolly, K. A. et al. A reservoir of stem-like CD8 T cells in the tumor-draining lymph node maintains the ongoing anti-tumor immune response. bioRxiv, 2021.2001.2027.428467, doi:10.1101/2021.01.27.428467 (2021).

41 Siddiqui, I. et al. Intratumoral Tcf1(+)PD-1(+)CD8(+) T Cells with Stem-like Properties Promote Tumor Control in Response to Vaccination and Checkpoint Blockade Immunotherapy. Immunity 50, 195–211.e110, doi:10.1016/j.immuni.2018.12.021 (2019).

42 Sekine, T. et al. TOX is expressed by exhausted and polyfunctional human effector memory CD8+ T cells. Science Immunology 5, eaba7918, doi:10.1126/sciimmunol.aba7918 (2020).

43 Alfei, F. et al. TOX reinforces the phenotype and longevity of exhausted T cells in chronic viral infection. Nature 571, 265–269, doi:10.1038/s41586-019-1326-9 (2019).

44 Khan, O. et al. TOX transcriptionally and epigenetically programs CD8+ T cell exhaustion. Nature 571, 211–218, doi:10.1038/s41586-019-1325-x (2019).

45 Scott, A. C. et al. TOX is a critical regulator of tumour-specific T cell differentiation. Nature 571, 270–274, doi:10.1038/s41586-019-1324-y (2019).

46 Rosato, P. C. et al. Virus-specific memory T cells populate tumors and can be repurposed for tumor immunotherapy. Nature Communications 10, 567, doi:10.1038/s41467-019-08534-1 (2019).

47 Shwetank et al. Maintenance of PD-1 on brain-resident memory CD8 T cells is antigen independent. Immunology and cell biology 95, 953–959, doi:10.1038/icb.2017.62 (2017).

48 Nayak, L., Lee, E. Q. & Wen, P. Y. Epidemiology of Brain Metastases. Current Oncology Reports 14, 48–54, doi:10.1007/s11912-011-0203-y (2012).

49 Ahmadzadeh, M. et al. Tumor antigen–specific CD8 T cells infiltrating the tumor express high levels of PD-1 and are functionally impaired. Blood 114, 1537–1544, doi:10.1182/blood-2008-12-195792 (2009).

50 Gros, A. et al. PD-1 identifies the patient-specific CD8+ tumor-reactive repertoire infiltrating human tumors. Journal of Clinical Investigation 124, 2246–2259, doi:10.1172/jci73639 (2014).

51 Gros, A. et al. Prospective identification of neoantigen-specific lymphocytes in the peripheral blood of melanoma patients. Nature Medicine 22, 433–438, doi:10.1038/nm.4051 (2016).

52 Lucca, L. E. et al. Circulating clonally expanded T cells reflect functions of tumor-infiltrating T cells. Journal of Experimental Medicine 218, doi:10.1084/jem.20200921 (2021).

53 Rosato, P. C. et al. Virus-specific memory T cells populate tumors and can be repurposed for tumor immunotherapy. Nature Communications 10, doi:10.1038/s41467-019-08534-1 (2019).

54 Simoni, Y. et al. Bystander CD8+ T cells are abundant and phenotypically distinct in human tumour infiltrates. Nature 557, 575–579, doi:10.1038/s41586-018-0130-2 (2018).

55 Shugay, M. et al. VDJdb: a curated database of T-cell receptor sequences with known antigen specificity. Nucleic Acids Research 46, D419–D427, doi:10.1093/nar/gkx760 (2018).

56 Hudson, W. H. & Sudmeier, L. J. Localization of T cell clonotypes using spatial transcriptomics. submitted.

57 Kaech, S. M. & Cui, W. Transcriptional control of effector and memory CD8+ T cell differentiation. Nature Reviews Immunology 12, 749–761, doi:10.1038/nri3307 (2012).

58 Voron, T. et al. VEGF-A modulates expression of inhibitory checkpoints on CD8+ T cells in tumors. Journal of Experimental Medicine 212, 139–148, doi:10.1084/jem.20140559 (2015).

59 Louveau, A. et al. CNS lymphatic drainage and neuroinflammation are regulated by meningeal lymphatic vasculature. Nature Neuroscience 21, 1380–1391, doi:10.1038/s41593-018-0227-9 (2018).

60 Hu, X. et al. Meningeal lymphatic vessels regulate brain tumor drainage and immunity. Cell Research 30, 229–243, doi:10.1038/s41422-020-0287-8 (2020).

61 Song, E. et al. VEGF-C-driven lymphatic drainage enables immunosurveillance of brain tumours. Nature 577, 689–694, doi:10.1038/s41586-019-1912-x (2020).

62 Lipson, E. J. et al. Relatlimab (RELA) plus nivolumab (NIVO) versus NIVO in first-line advanced melanoma: Primary phase III results from RELATIVITY-047 (CA224-047). Journal of Clinical Oncology 39, 9503–9503, doi:10.1200/JCO.2021.39.15_suppl.9503 (2021).

63 Tawbi, H. A. et al. Combined Nivolumab and Ipilimumab in Melanoma Metastatic to the Brain. N Engl J Med 379, 722–730, doi:10.1056/NEJMoa1805453 (2018).

64 Van Gassen, S. et al. FlowSOM: Using self-organizing maps for visualization and interpretation of cytometry data. Cytometry Part A 87, 636–645, doi:10.1002/cyto.a.22625 (2015).

65 Mclnnes, L., Healy, J. & Melville, J. UMAP: Uniform Manifold Approximation and Projection for Dimension Reduction. arXiv pre-print server, doi:None arxiv:1802.03426 (2020).

66 Stuart, T. et al. Comprehensive Integration of Single-Cell Data. Cell 177, 1888–1902.e1821, doi:10.1016/j.cell.2019.05.031 (2019).

67 Wickham, H. in Use R!, 1 online resource (XVI, 260 pages 232 illustrations, 140 illustrations in color (Springer International Publishing : Imprint: Springer, Cham, 2016).

68 Korotkevich, G. et al. Fast gene set enrichment analysis. biorXiv, doi:10.1101/060012 (2021).

69 Sadee, C., Pietrzak, M., Seweryn, M., Wang, C. & Rempala, G. divo: Tools for Analysis of Diversity and Similarity in Biological Systems, <https://CRAN.R-project.org/package=divo> (2019).

70 Oksanen, J. et al. vegan: Community Ecology Package, <https://CRAN.R-project.org/package=vegan> (2020).

71 Bagaev, D. V. et al. VDJdb in 2019: database extension, new analysis infrastructure and a T-cell receptor motif compendium. Nucleic Acids Research 48, D1057–D1062, doi:10.1093/nar/gkz874 (2020).

72 Robins, H. S. et al. Comprehensive assessment of T-cell receptor β-chain diversity in αβ T cells. Blood 114, 4099–4107, doi:10.1182/blood-2009-04-217604 (2009).

73 Bolotin, D. A. et al. MiXCR: software for comprehensive adaptive immunity profiling. Nature Methods 12, 380–381, doi:10.1038/nmeth.3364 (2015).

