## Supplementary Information for "The CD8^+^ T cell landscape of human brain metastases"

**Supplementary Table 1: Tumor Histology**

| Primary Tumor Histology | n |
|---|---|
| Lung carcinoma (4 SCLC, 10 NSCLC) | 14 |
| Breast carcinoma | 6 |
| Melanoma | 4 |
| Esophageal carcinoma | 1 |
| Poorly differentiated carcinoma, unspecified primary | 2 |
| Adenoid cystic carcinoma | 1 |
| Uterine carcinoma | 1 |
| Renal cell carcinoma | 1 |
| Urothelial carcinoma | 1 |

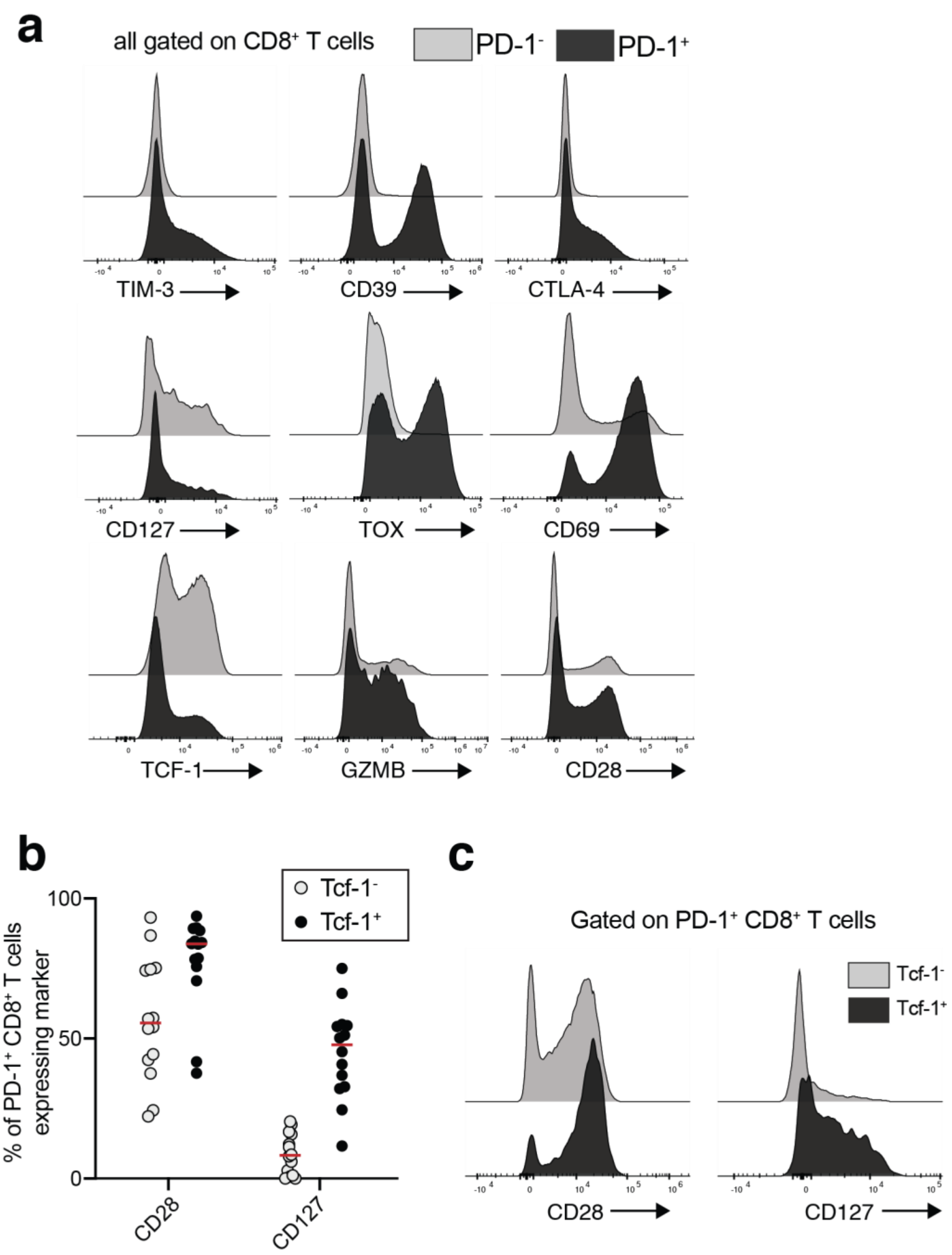


**Supplementary Figure 1: Flow cytometry analysis of brain metastasis-infiltrating CD8$^+$ T cells** (a) Representative flow plots for markers shown in Figure 1f. (b) Comparison of CD28 and CD127 expression on TCF-1$^-$ and TCF-1$^+$ PD-1$^+$ CD8$^+$ T cells. Bars on graph indicate medians. (c) Representative flow plots of each marker in (b).

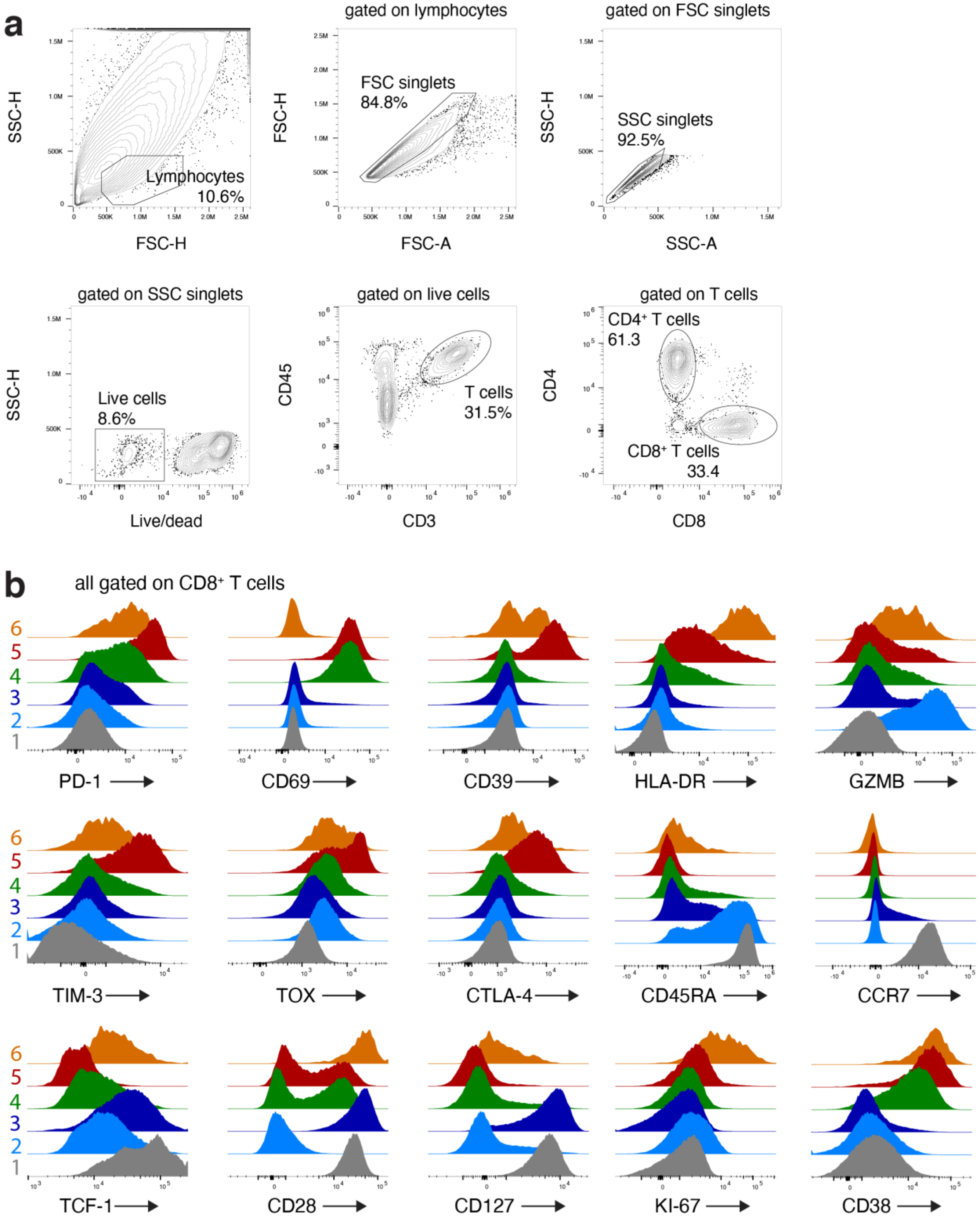


**Supplementary Figure 2: High-parameter flow cytometry analysis of CD8⁺ T cells.** (a) Gating strategy for CD8⁺ T cells. (b) Expression of all markers used for UMAP in each CD8⁺ T cell cluster.

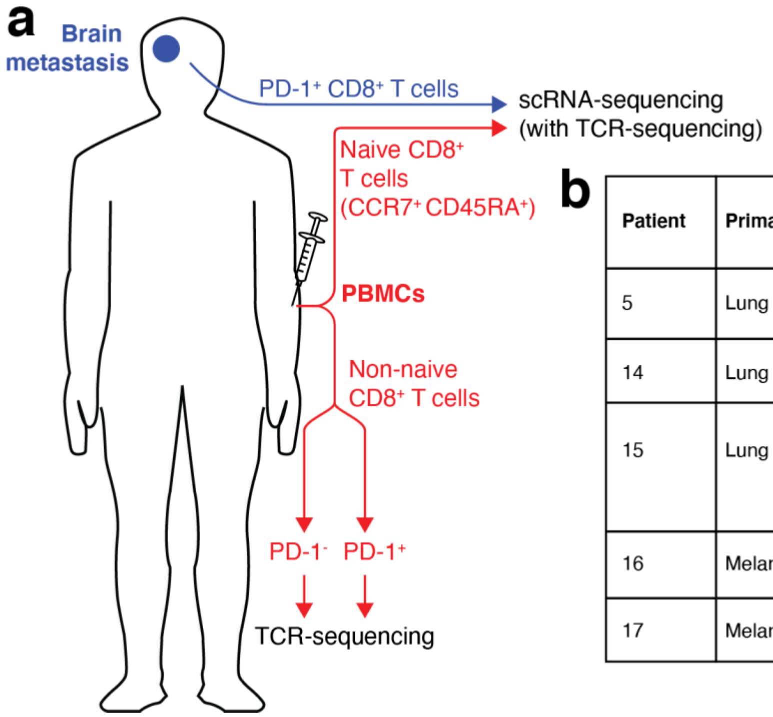


**b**

| Patient | Primary | Lymphocytes/ g of tumor | CD8+ T cells/ g of tumor | PD-1+ (% of CD8+) |
|---|---|---|---|---|
| 5 | Lung | $1.8 \times 10^6$ | $3.3 \times 10^5$ | 86 |
| 14 | Lung | $1.1 \times 10^6$ | $2.3 \times 10^5$ | 73 |
| 15 | Lung | $1.2 \times 10^6$ | $3.8 \times 10^5$ | 70 |
| 16 | Melanoma | $4.9 \times 10^6$ | $2.3 \times 10^6$ | 74 |
| 17 | Melanoma | $2.1 \times 10^7$ | $1.5 \times 10^7$ | 95 |

**Supplementary Figure 3**: **Schematic and patient data for single cell RNA-sequencing experiments.** a) PD-1$^+$ CD8$^+$ T cells were isolated from brain metastases by FACS immediately after surgical resection. These were mixed with naïve CD8$^+$ T cells from peripheral blood after staining with hashing antibodies and subjected to scRNA-seq. Non-naïve CD8$^+$ T cells from blood were sorted into PD-1$^-$ and PD-1$^+$ populations and separately subjected to TCRβ sequencing. (b) Patient data for scRNA-seq samples. All lung primaries are non-small cell lung tumors.

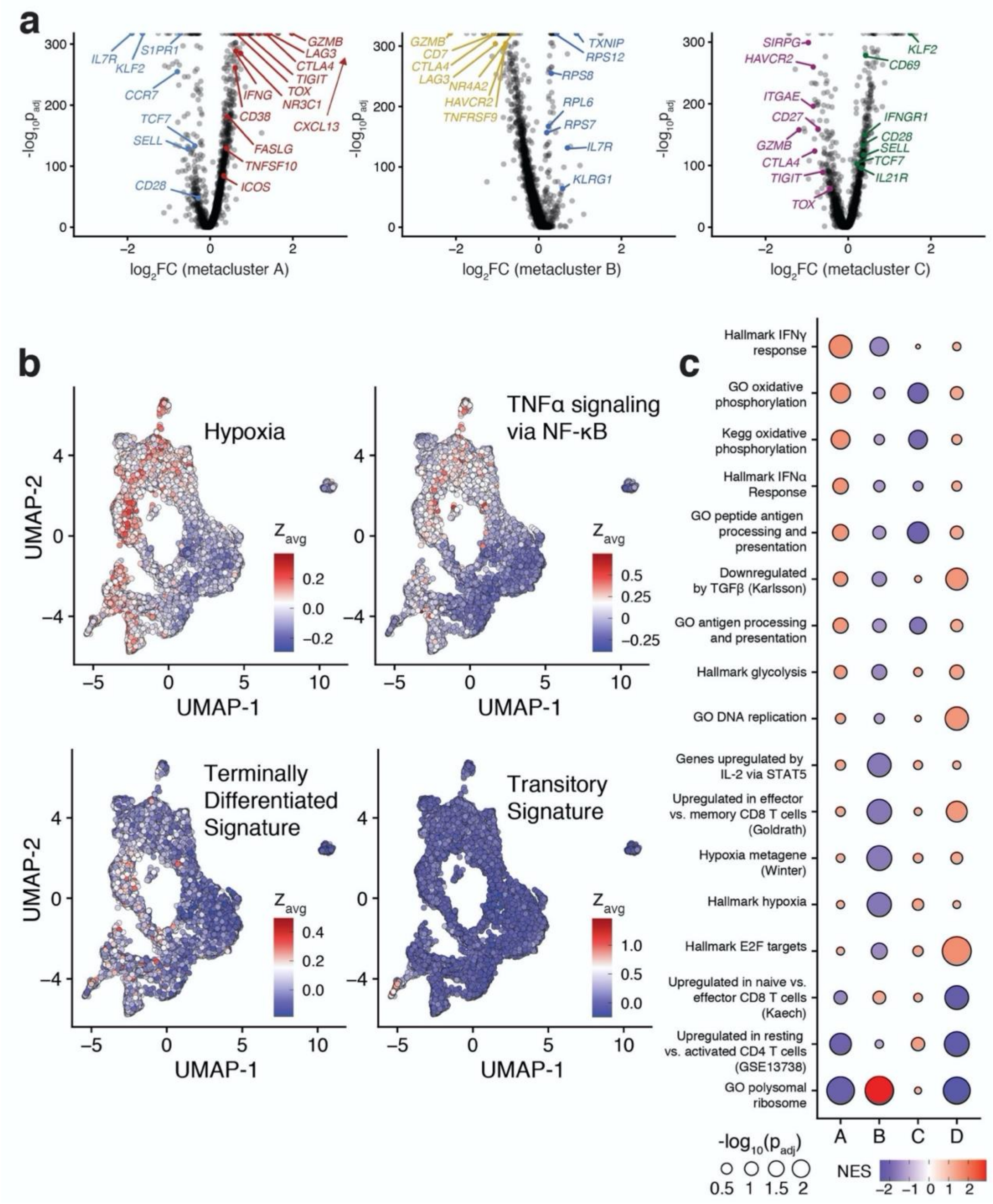


**Supplementary Figure 4: The transcriptional signature of transitory cells from the murine LCMV system is enriched in the Dividing metacluster.** (a) Volcano plots showing selected highly differentially-expressed genes in each metacluster compared with all other non-naïve metaclusters. (b) Expression of selected gene sets in each cell, projected on the UMAP. The terminally differentiated and transitory gene sets are based on the transcriptional phenotype of CD8$^{+}$ T cells in the murine LCMV system. Cells are colored by the median z-score of genes in the indicated gene set. (c) Net enrichment score and statistical significance of selected gene sets in each cluster.

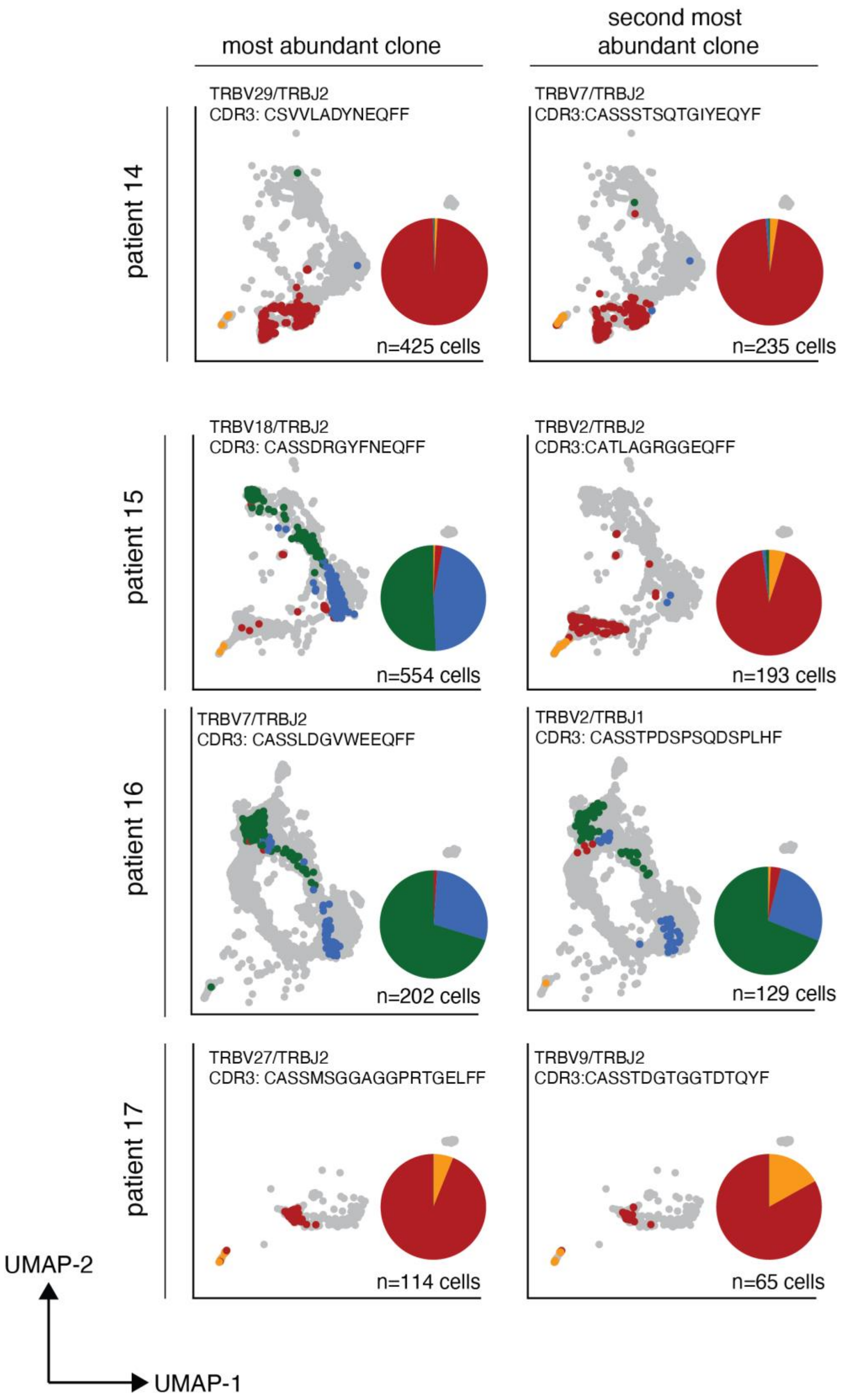


**Supplementary Figure 5: Cells within the most abundant clones of brain metastasis-infiltrating PD-1$^+$ CD8$^+$ T cells exhibit restriction to a metacluster A/D or B/C phenotype.** Phenotype of the most abundant (left) and second most abundant (right) TCR clones from each scRNA-seq sample. Patient 5 is shown in Figure 4f. Cells from other clones of the indicated patient are shown in gray. Inlaid pie charts show distribution of cells by metacluster for the indicated clone.

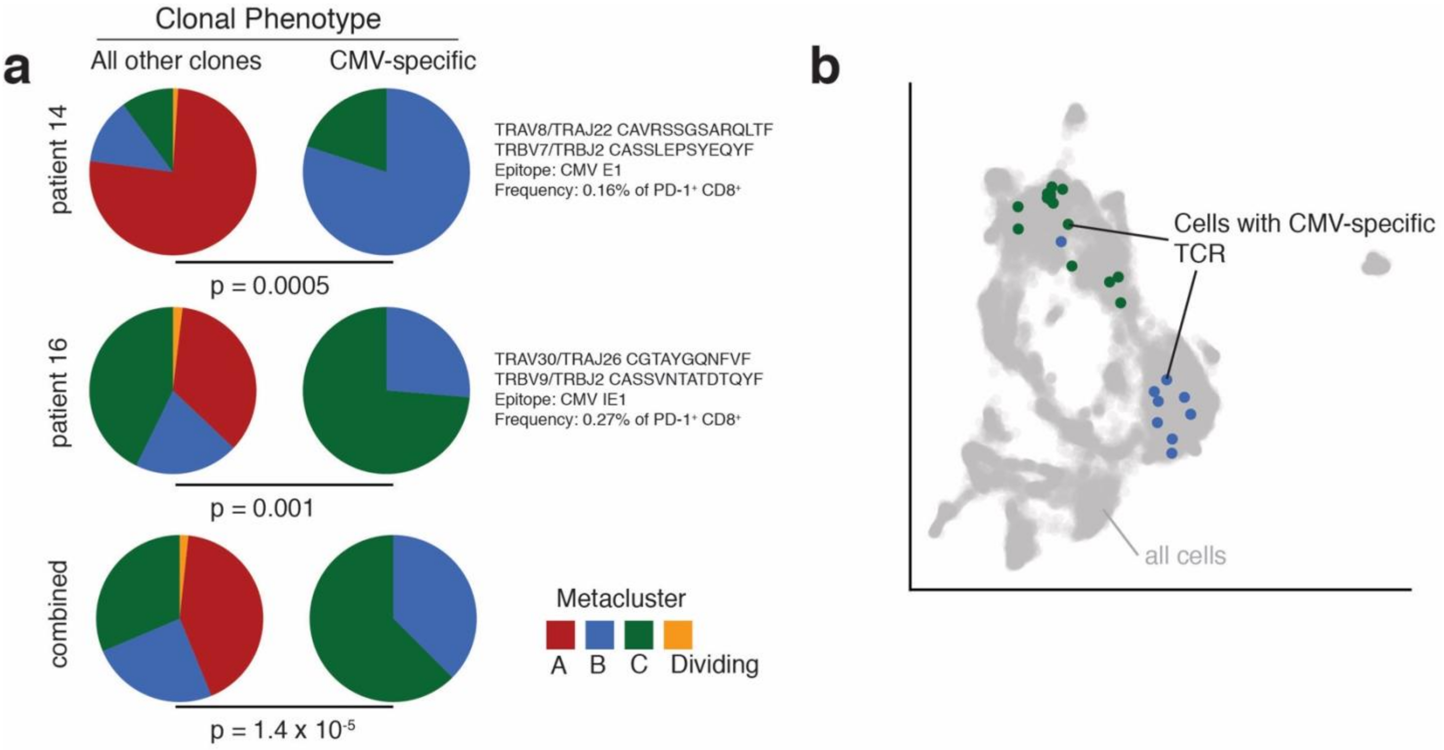


**Supplementary Figure 6: Phenotype of brain metastasis-infiltrating PD-1$^+$ CD8$^+$ T cells expressing CMV-specific TCRs identified in the VDJdb.** (a) One CMV-specific TCR was found in patient 14, and one was found in patient 16. Pie charts indicate the scRNA-seq phenotype of CMV-specific clones and all other clones among brain-metastasis PD-1$^+$ CD8$^+$ T cells. (b) UMAP with brain metastasis-infiltrating PD-1$^+$ CD8$^+$ T cells expressing a CMV-specific TCR identified in the VDJdb, colored by phenotype. P-values in (a) were calculated with Fisher's exact test.

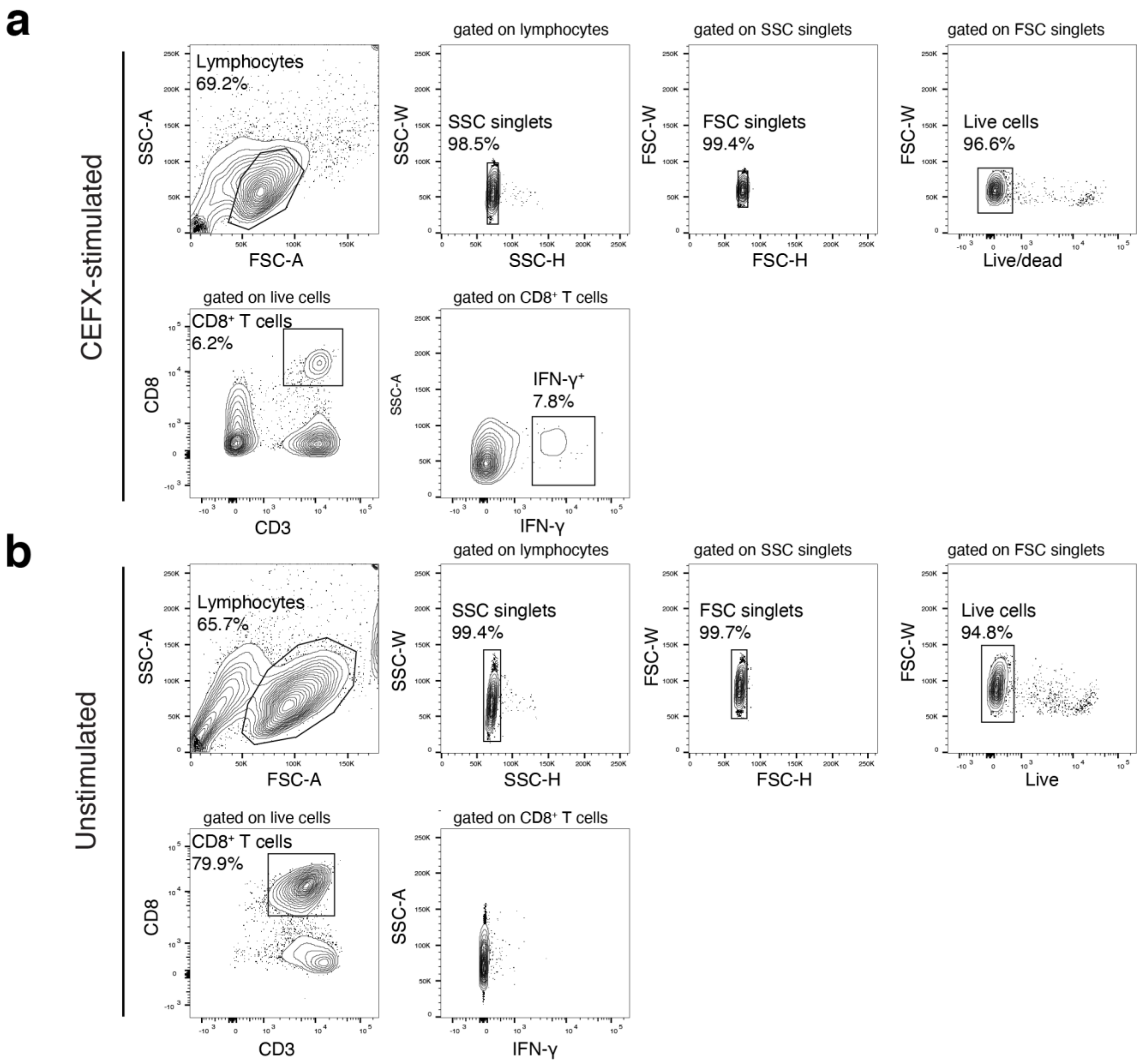


**Supplementary Figure 7: Gating strategy for IFN-γ capture assay to determine CEFX-reactive CD8⁺ T cells.** (a) Expanded and CEFX-stimulated PBMCs from a brain metastasis patient. (b) Expanded, unstimulated PBMCs from a healthy donor.

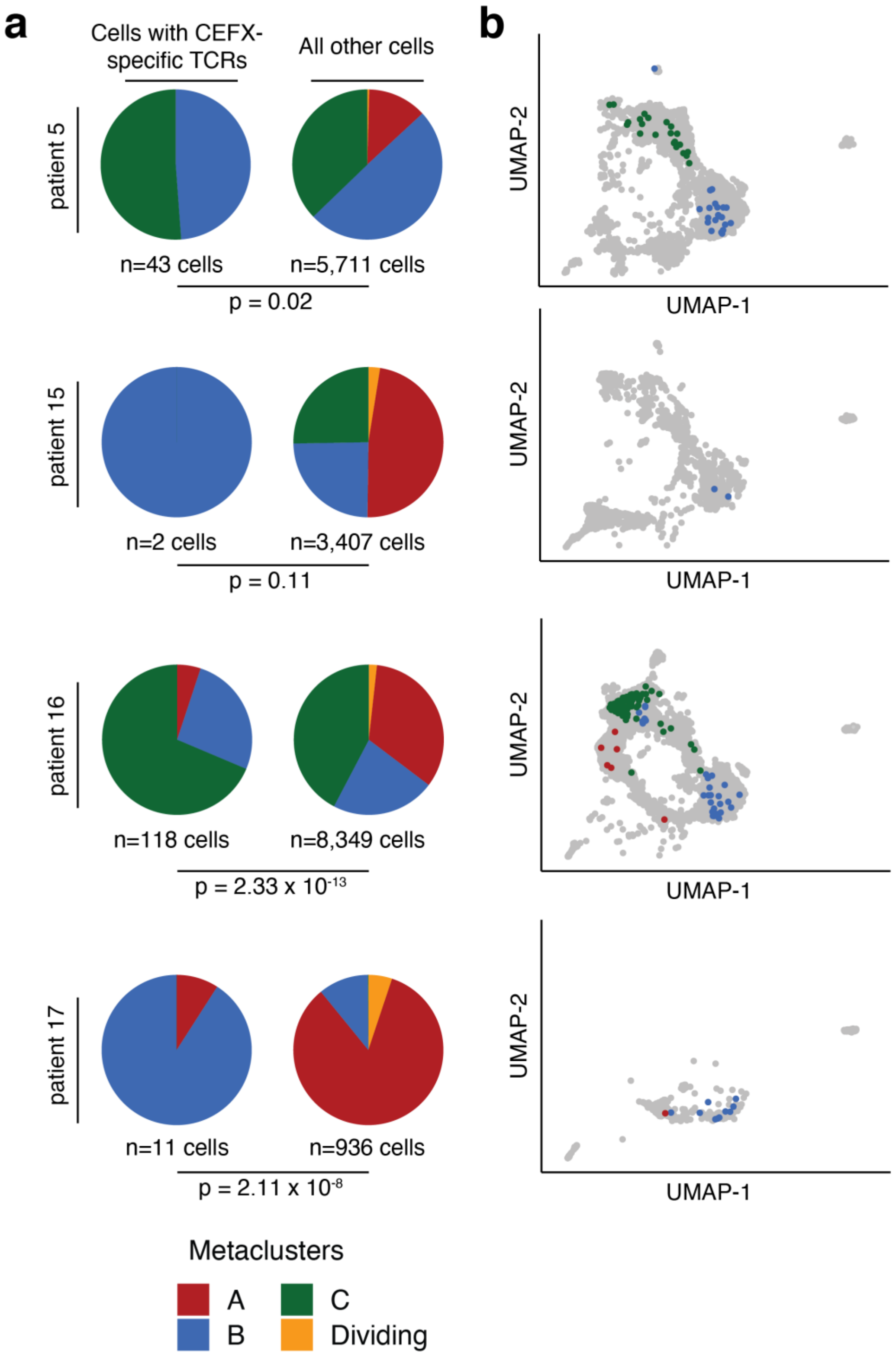


**Supplementary Figure 8: Phenotype of brain metastasis-infiltrating CEFX-specific PD-1⁺ CD8⁺ T cells from each patient.** PBMCs were not available from patient 14 for this experiment. CEFX-specific CD8⁺ T cells were determined by their expression of TCRs identified in the IFN-γ capture assay shown in Figure 5. (a) Distribution of CEFX-specific (left) and all other (right) brain metastasis-infiltrating PD-1⁺ CD8⁺ T cells by scRNA-seq metacluster. (b) UMAP of brain metastasis-infiltrating PD-1⁺ CD8⁺ T cells, with CEFX-specific cells colored by metacluster. All other cells from the patient are shown in gray. P-values in (a) were calculated with Fisher's Exact Test.

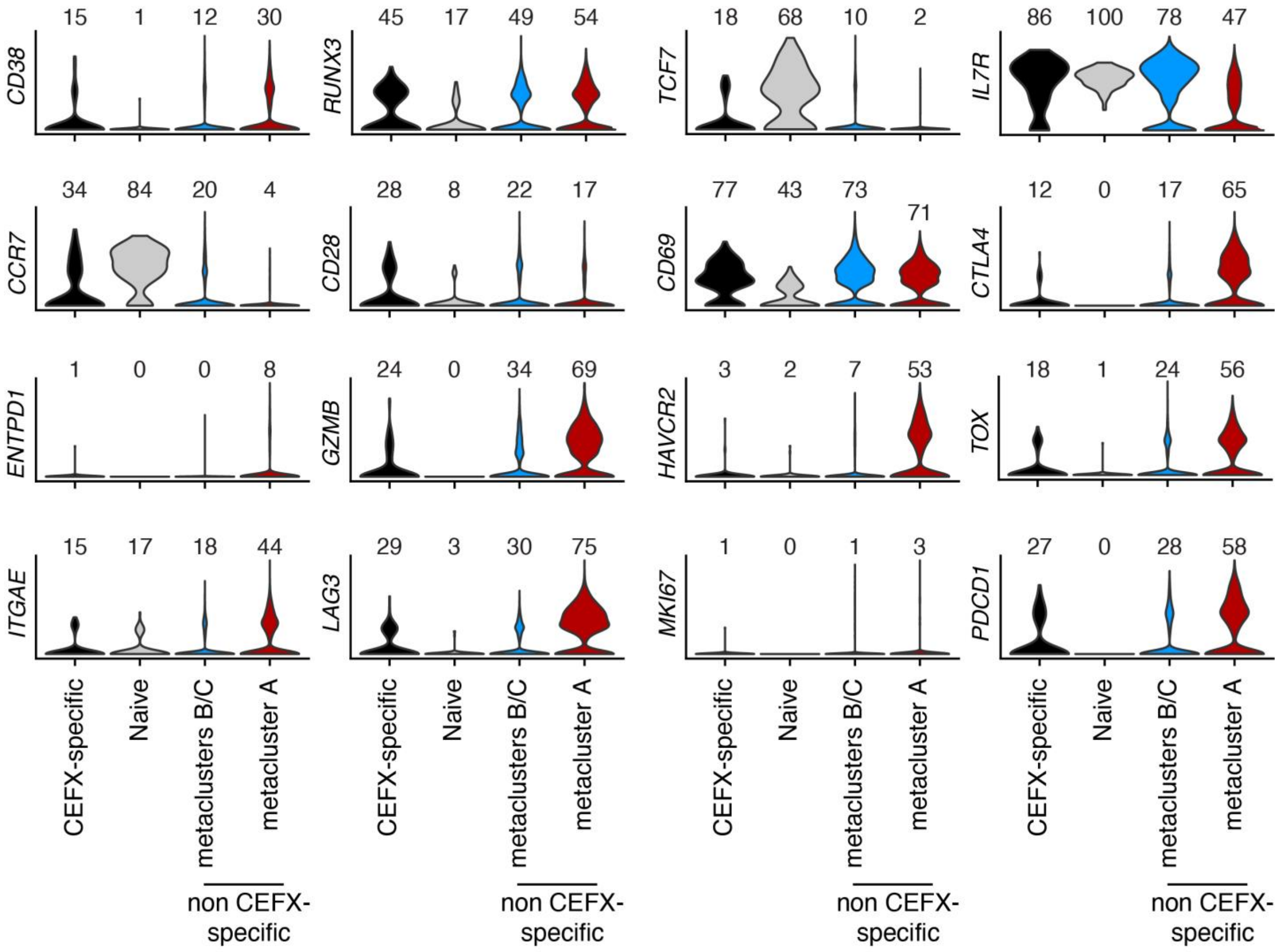


**Supplementary Figure 9: Expression of selected genes by CEFX-specific and other brain metastasis-infiltrating PD-1⁺ CD8⁺ T cells**. The number above each column indicates percent of cells with measurable expression of each gene.

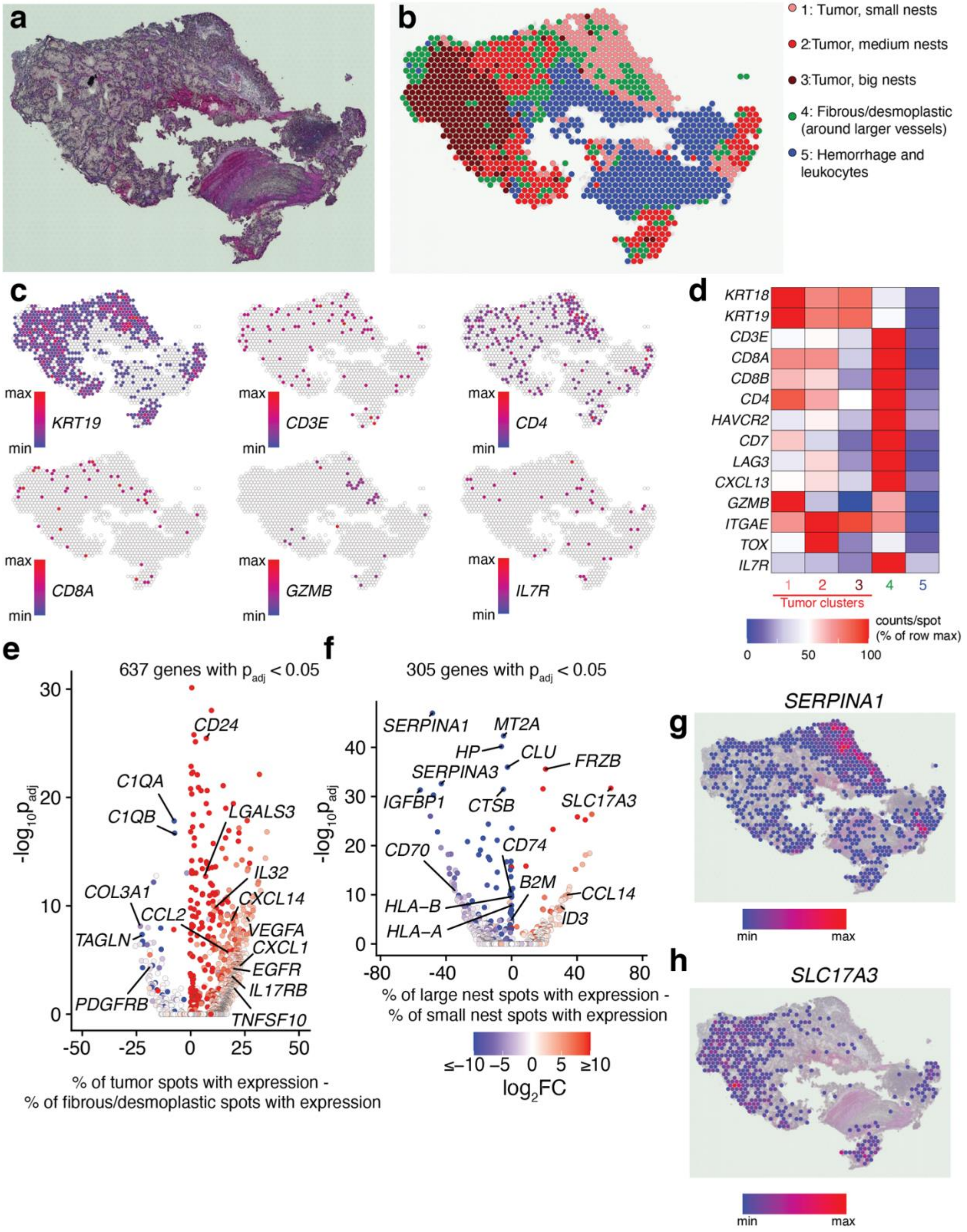


**Supplementary Figure 10: A renal clear cell brain metastasis exhibits transcriptional heterogeneity based on tumor nest size.** (a) Hematoxylin and eosin-stained (H&E) section of a renal clear cell brain metastasis (patient 24). (b) Graph-based clusters of gene expression with annotations. (c) Spatial expression of selected genes. (d) Heatmap of normalized gene expression density in each cluster. (e) Differential gene expression between tumor (clusters 1, 2, and 3) and surrounding fibrous/desmoplastic tissue (cluster 4). (f) Differential gene expression between large nests of tumor (cluster 3) and small nests of tumor (cluster 1). (g-h) Expression of two example genes highly expressed in small nests (*SERPINA1*, panel g) and large nests (*SLC17A3*, panel h).

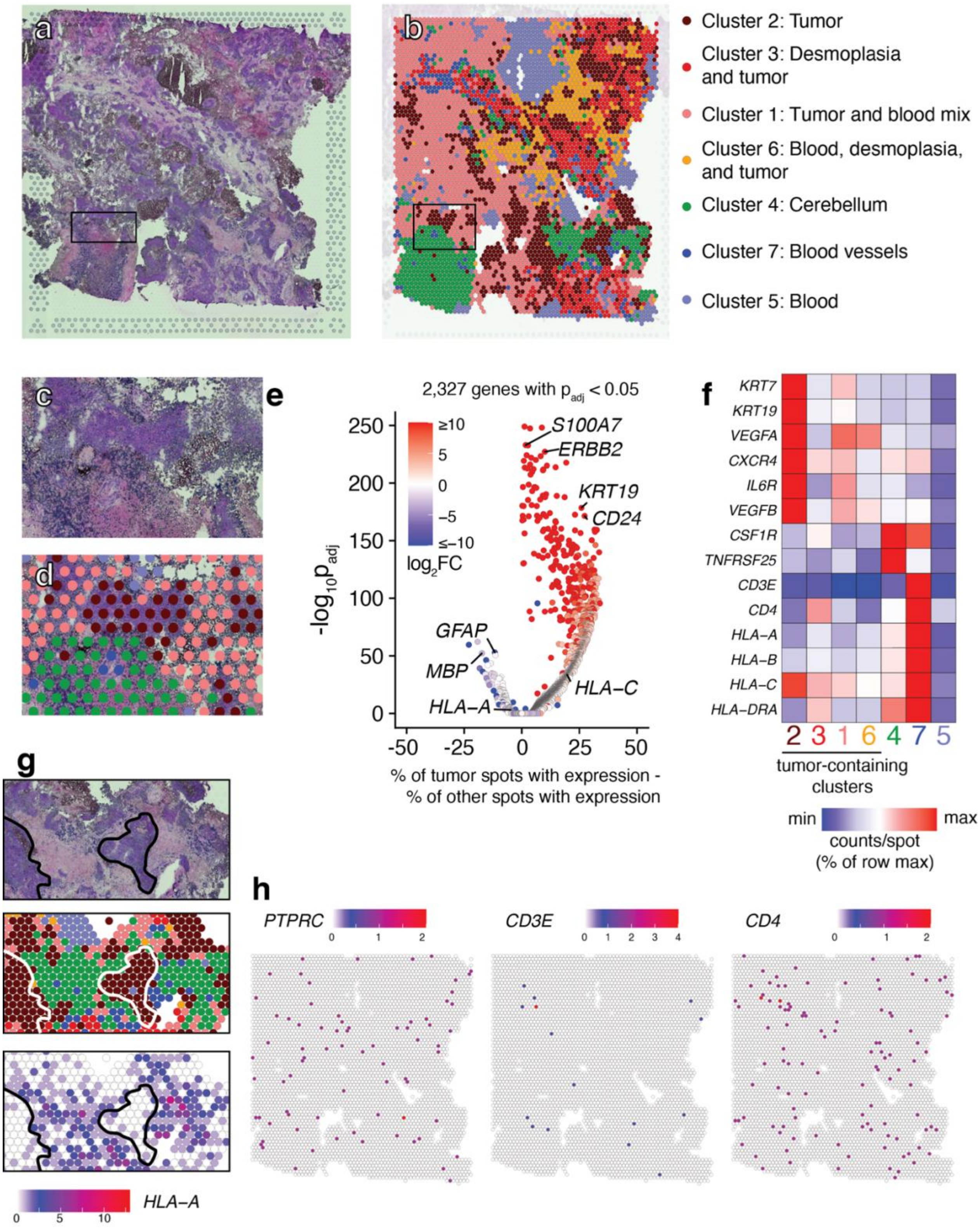


**Supplementary Figure 11: Spatial transcriptomics of a poorly-infiltrated brain metastasis of breast carcinoma reveals spatial heterogeneity of gene expression and loss of MHC I genes by tumor regions.** a) H&E-stained section of a breast carcinoma brain metastasis (patient 26). b) Annotated graph-based clusters of gene expression. c-d) Inset of H&E staining and clusters. e) Differential gene expression of tumor-containing regions (clusters 1, 2, 3, and 6) compared to other clusters. f) Heatmap of normalized gene expression density in each cluster. g) Loss of *HLA-A* expression in tumor clusters. Top panel: H&E staining; middle: graph-based clusters; bottom panel: expression of *HLA-A*. Selected tumor regions are outlined. h) Expression of *PTPRC* (encoding CD45), *CD3E*, a pan T cell marker, and *CD4*. *CD8A* expression was below the limit of detection.

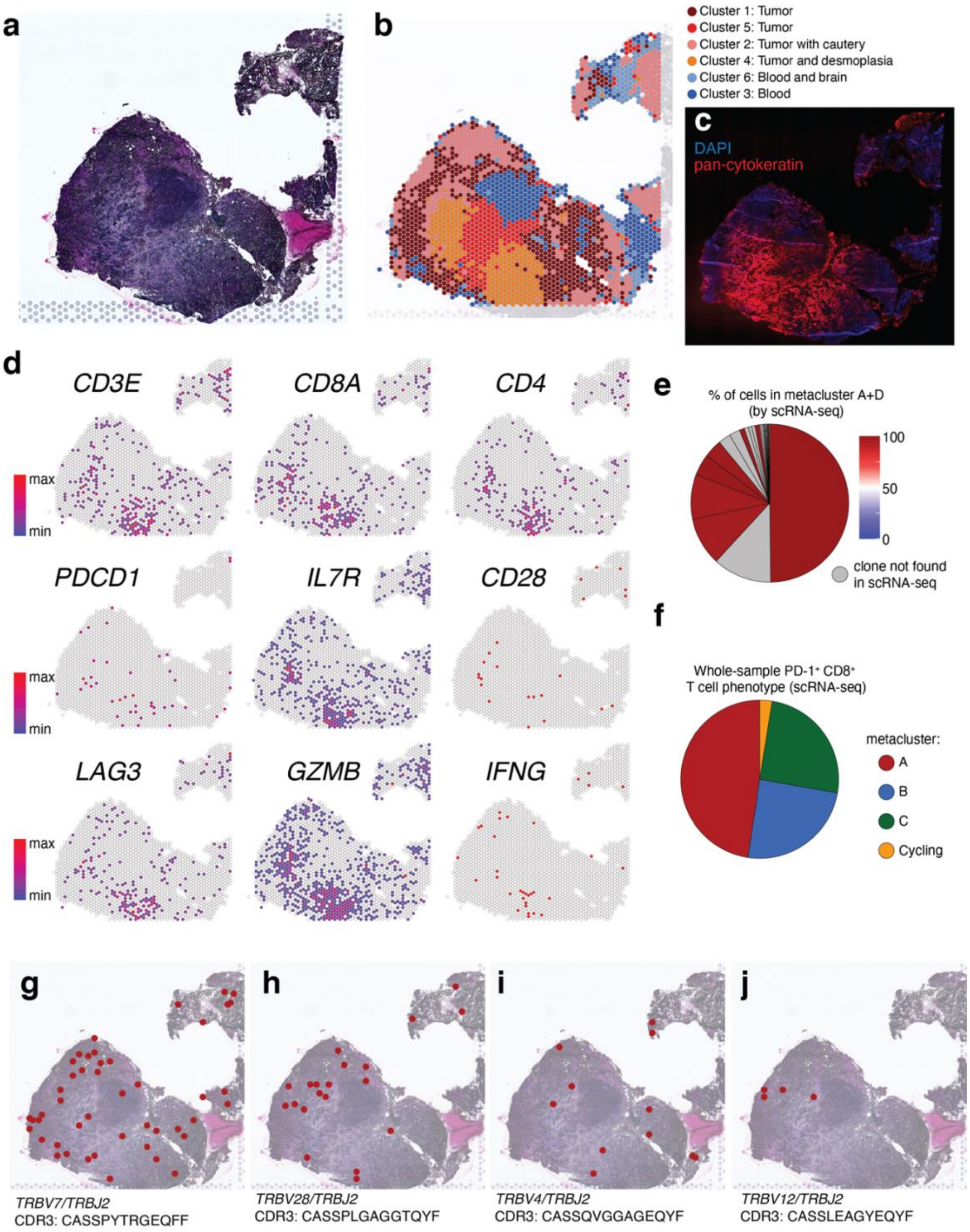


**Supplementary Figure 12: Spatial gene expression and TCR sequencing of a lung adenocarcinoma brain metastasis (patient 15).** a) H&E staining of the metastasis. b) Gene expression clusters identified by graph-based clustering. c) Immunofluoresence of an adjacent section with DAPI and pan-cytokeratin (a tumor marker) demonstrates this tissue section is comprised entirely of tumor parenchyma. d) Spatial expression of selected immune-related genes. e) Spatial TCR sequencing was performed on this tumor section. Shown are the T cell clones identified within this section, colored by their scRNA-seq phenotype, if applicable. Size the of pie chart slice indicates TCR frequency among spatially-identified clones. All clones found both in scRNA-seq and spatial TCR sequencing were composed of terminally-differentiated cells from metaclusters A and D. f) scRNA-seq metacluster of all cells sequenced from the matched patient sample. When compared to panel (e), terminally differentiated cells were highly enriched in the tumor parenchyma. (g-j) Spatial distribution of four example TCR clones throughout the tissue section.

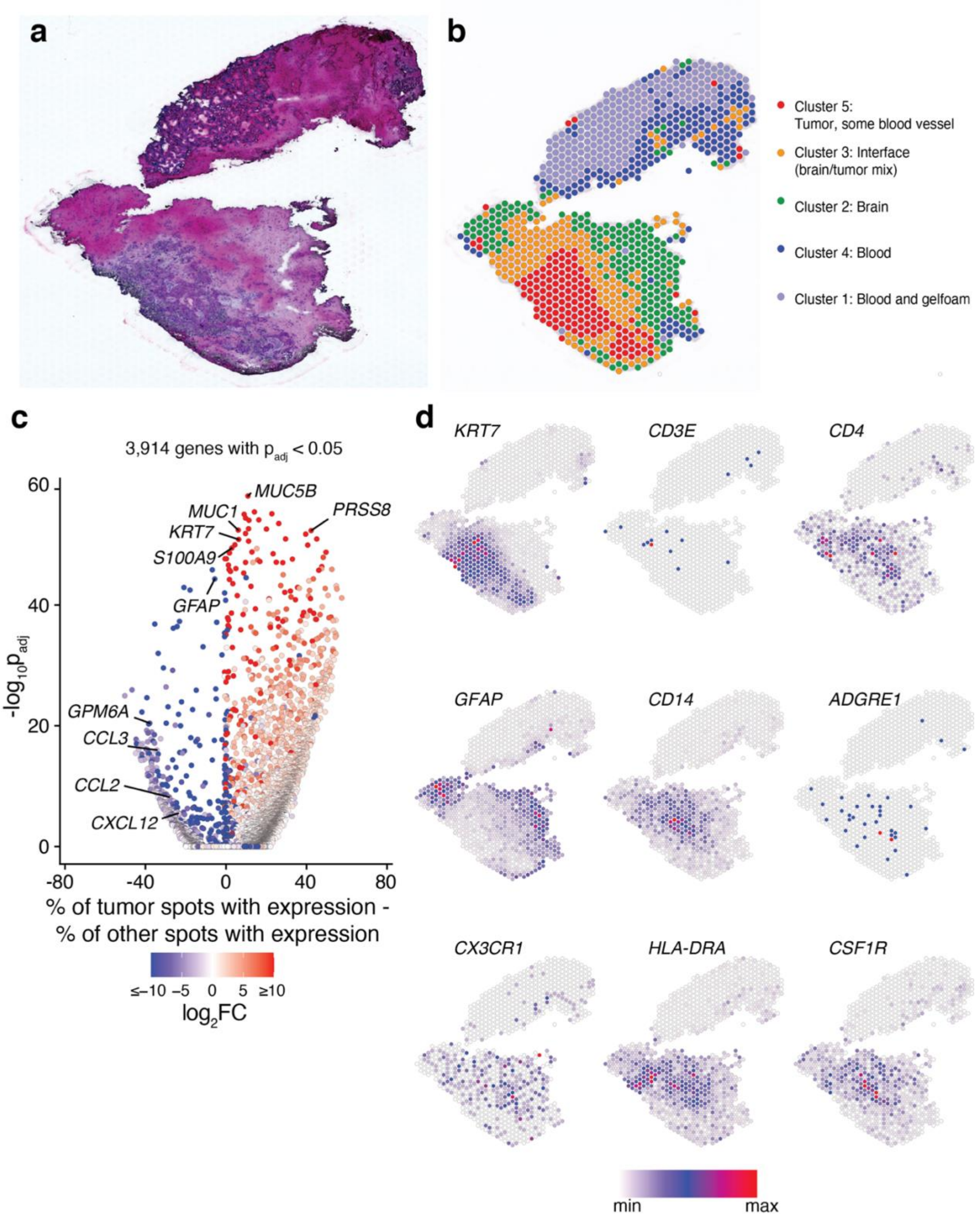


**Supplementary Figure 13: Spatial gene expression of a lung adenocarcinoma brain metastasis tissue section poorly infiltrated by CD8⁺ T cells (patient 19).** a) H&E staining of a lung adenocarcinoma brain metastasis. b) Spatial distribution of annotated gene expression clusters. c) Differences in gene expression between capture spots with tumor (cluster 5) and all other clusters. d) Spatial expression of selected genes shows high expression of *KRT7*, a cytokeratin protein, within the tumor. T cell infiltrate (*CD3E*) is sparse and located at the brain/tumor interface, but high levels of innate immune-associated genes such as *CD14* are also found around the tumor. *GFAP*, expressed in glial cells such as astrocytes, is expressed beyond the tumor/brain interface.

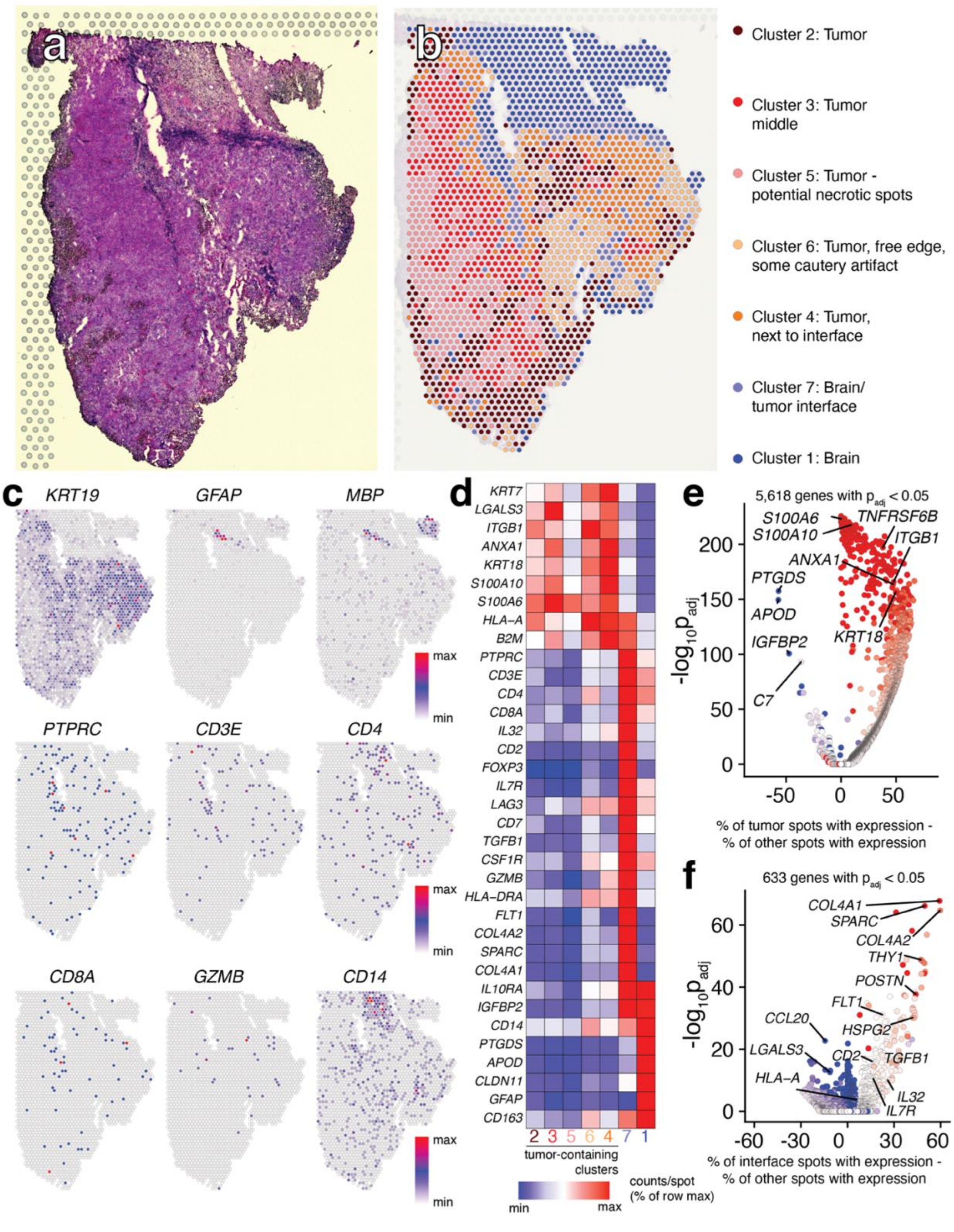


**Supplementary Figure 14: Spatial gene expression within a lung adenocarcinoma tissue section (patient 26).** a) H&E staining of a lung adenocarcinoma brain metastasis. b) Spatial distribution of annotated gene expression clusters. c) Expression of selected genes, including *KRT19* (a cytokeratin and tumor marker), *GFAP*, and *MBP* (myelin basic protein, expressed by Schwann cells and oligodendrocytes). d) Heatmap of selected gene expression density by cluster. e) Volcano plot of gene expression differences in tumor spots (clusters 2, 3, 4, 5, and 6) compared to other tissue spots. f) Gene expression differences at the brain/tumor interface (cluster 7) compared to all other spots.

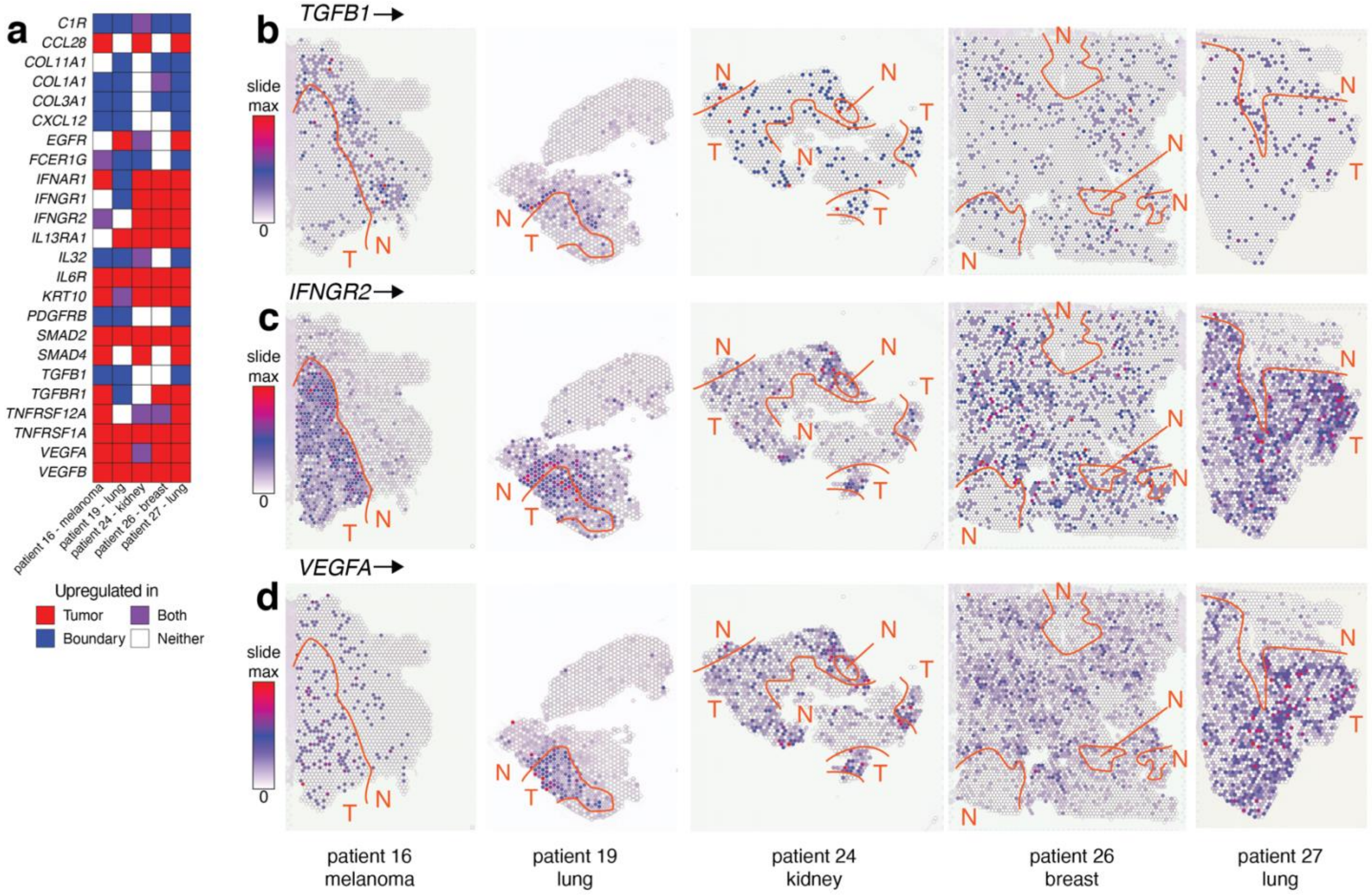


**Supplementary Figure 15: Spatial location of cytokine, chemokine, and their receptor genes in human brain metastases.** a) Heatmap showing sites of selected gene upregulation in five brain metastases samples. Sample 15 was excluded from analysis as it contained only tumor parenchyma with no margin. Upregulation is defined as higher expression and an adjusted p-value of <0.05 in the tumor, boundary, or both compared to all other tissue spots. b-d) Spatial expression of *TGFB1* (encoding TGF-β), *IFNGR2* (encoding the β chain of the IFN-γ receptor), and *VEGFA*, which encodes the Vascular Endothelial Growth Factor A. Tissue annotations for approximate regions of tumor (T) or non-tumor (N) are shown. See Fig. 6 and Supplementary Figs. 10, 11, 13, and 14 for more detailed tissue annotation.

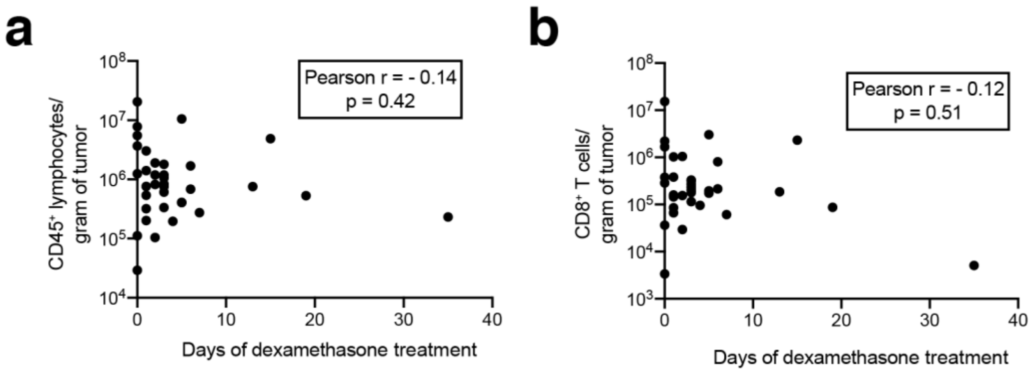


**Supplementary Figure 16: Days of dexamethasone therapy is not correlated with density of immune cell infiltration in brain metastases.** Days of treatment with dexamethasone versus immune cell infiltration are shown for $CD45^+$ lymphocytes (a) and $CD8^+$ T cells (b).
